# Warming and precipitation change alter flowering phenology and coflowering networks in California serpentine grasslands

**DOI:** 10.64898/2026.08.05.743098

**Authors:** Andrea N. Nebhut, Jeffrey S. Dukes

**Author notes:** Author for correspondence: Andrea Nebhut.

## Abstract

Coflowering, the temporal overlap in flowering among plant species, can influence plant fitness through its effects on heterospecific pollen transfer and competition for pollinators and resources. This temporal overlap is changing as plant species respond differently to climate change. However, difficulties in quantifying and comparing patterns of coflowering across diverse, multispecies communities have limited progress in understanding climate-driven shifts in coflowering. Network-based approaches offer a promising solution to this limitation. Here, we investigate how warming and altered precipitation influence coflowering network structure, flowering phenology, and seed production in 12 annual California serpentine grassland plant species in a mesocosm experiment, using coflowering network analysis and structural equation modeling. Communities consistently grouped into two phenological modules corresponding to early- and late-season annuals. Warming reduced the overall amount of coflowering in the community, driven primarily by weakened and less diverse coflowering among late-season species. Soil moisture moderated these effects: late-season coflowering was strongest under cool–wet conditions and weakest under warm conditions regardless of soil moisture. In contrast, early-season species had a more fixed phenological window, and maintained stable coflowering relationships across climates. Growth form (grass or forb) and origin (native vs. non-native) did not predict coflowering responses. We additionally found that temperature and soil moisture influenced seed production largely through their effects on flowering phenology, with early- and late-season species exhibiting distinct phenological responses and sensitivities. Our results demonstrate that warming and precipitation change can spread out flowering times within functional groups, reshaping community-wide coflowering networks through species-specific responses to altered climate conditions.

## Introduction

Phenology – the timing of recurring life-history events – is sensitive to climatic variation and change (Menzel et al. 2006; Jeong et al. 2011; Gill et al. 2015; Thackeray et al. 2016; Cohen et al. 2018; Piao et al. 2019; Romano et al. 2022; Ramirez-Parada et al. 2024). Among plants, the onset, peak, and cessation of flowering represent particularly consequential developmental transitions governed by cues such as photoperiod, temperature, and soil moisture (Rathcke and Lacey 1985; Sherry et al. 2007). Warming generally advances or extends flowering (Cleland et al. 2006; Menzel et al. 2006; Sherry et al. 2007; Ellwood et al. 2013; CaraDonna et al. 2014; Whittington et al. 2015; Valencia et al. 2016; Moore and Lauenroth 2017), but responses are not uniform; warming can instead delay or shorten flowering in some species or produce no detectable phenological change in others (Fitter et al. 1995; Fitter and Fitter 2002; Haiying et al. 2010). Research on soil moisture effects is more limited, but similarly heterogeneous. Decreased water availability can lead to earlier (Llorens and Peñuelas 2004; Jentsch et al. 2009; Gordo and Sanz 2010; Lesica and Kittelson 2010; Moore and Lauenroth 2017), later (Peñuelas et al. 2004; Nagy et al. 2013), unchanged (Sparks and Carey 1995; Thórhallsdóttir 1998; Abu-Asab et al. 2001; Bloor et al. 2010), or functional-group-dependent flowering responses (Castillioni et al. 2022). When warming and drying occur simultaneously, their combined effects can intensify phenological shifts beyond those caused by either factor alone (Bloor et al. 2010; Gugger et al. 2015; Matthews and Mazer 2015; Zhou et al. 2023), but synergistic effects are not universal (Arfin Khan et al. 2018). Consequently, phenological outcomes are strongly species- and context-dependent (Pérez-Ramos et al. 2020), and because plants may shift the onset, peak, and end of flowering independently, evaluation of multiple flowering phases is necessary for understanding climate-phenology relationships (Nagy et al. 2013; CaraDonna et al. 2014; Jiang et al. 2016).

Changes in flowering phenology can affect plant fitness. Individuals that flower before accumulating adequate resources may suffer reduced seed production, while delayed flowering risks the successful completion of seed production before the end of the season (Elzinga et al. 2007). Furthermore, mismatches between individual and population-level phenology can reduce reproductive success because pollination visitation rates typically peak with community-wide floral abundance, and seeds set during peak bloom may escape predation through predator satiation, whereas those produced earlier or later incur greater predation risks (Elzinga et al. 2007). Phenological change can also alter species interactions, with potential fitness consequences. Much of the recent literature about the effects of phenological change on species interactions has focused on “mismatches” across trophic levels when interacting species shift their phenology at different rates, particularly mismatches between plants and their pollinators or herbivores (Tylianakis et al. 2008; Renner and Zohner 2018; Kharouba et al. 2018). However, phenological restructuring within a trophic level also has consequences for species interactions and fitness. Increases in coflowering, the amount of temporal overlap among plant species (Arceo-Gómez et al. 2018), may enhance pollination success by attracting more and a greater diversity of pollinators (Ghazoul et al. 2006; Liao et al. 2011), but may also increase interspecific pollinator competition and heterospecific pollen transfer, reducing seed set (Mitchell et al. 2009; Mesgaran et al. 2017; Tiusanen et al. 2020). Greater coflowering can also intensify competition for shared soil resources by causing greater convergence in the times of peak resource use between neighboring plants (Gulmon et al. 1983; Nord and Lynch 2009; Cleland and Wolkovich 2024).

Coflowering can be altered when plant species differ in their phenological responses to changing climate conditions (Aldridge et al. 2011; Fisogni et al. 2022), but predicting community-level phenological responses to climate change remains challenging. First, species’ phenological responses to warming and drying are highly variable, leading different communities to diverge or converge in phenological distributions under the changing climate (Diez et al. 2012). Even when species respond in a consistent direction (e.g., all advance flowering), subtle interspecific differences in the timing, duration, or intensity of flowering can shift coflowering patterns (Forrest et al. 2010). However, some trends have emerged that suggest that functional traits and evolutionary histories may structure phenological responsiveness. Previous work suggests that early-flowering species are often more responsive than late bloomers (Menzel et al. 2006; Sherry et al. 2007; Wolkovich et al. 2012; Calinger et al. 2013), forbs respond more strongly than grasses (Iverson et al. 2009), and non-native species tend to be more responsive than native species (Lesica and Kittelson 2010; Calinger et al. 2013; Zettlemoyer et al. 2019). Second, the difficulty of quantifying and comparing coflowering dynamics across diverse, multispecies communities has limited such analyses to examining differences in coflowering among only a small number of species (Forrest et al. 2010), aggregating flowers across species to assess community-wide trends in flower abundance (Aldridge et al. 2011), comparing patterns in the strength of coflowering for pairs of species within a diverse community (Fisogni et al. 2022; Pareja-Bonilla et al. 2025), or comparing differences in the composition (Ramirez-Parada et al. 2025) or richness (Theobald et al. 2017; Park et al. 2025) of coflowering communities.

Network analysis of coflowering allows us to quantify patterns of community-level flowering overlap, examine the strength and diversity of coflowering relationships experienced by single species, and identify groups of species that flower together and the species that bridge these phenological modules (Arceo-Gómez et al. 2018). Prior work with coflowering networks has focused primarily on evolutionary and pollination-driven explanations for coflowering network structure (Arceo-Gómez et al. 2018; Albor et al. 2020; Albor et al. 2022; Suárez-Mariño 2022a;

Suárez-Mariño 2022b), and this approach has rarely been applied to global change contexts; the only example to date shows that non-target herbicide exposure can reduce coflowering strength and restructure phenological modules (Iriart et al. 2022). To our knowledge, no study has evaluated how climate change–induced warming and precipitation change may reshape coflowering networks. Therefore, in this study, we assessed how warming and altered precipitation influenced coflowering, flowering phenology, and seed production in California serpentine grassland communities. Specifically, we asked:

1. *How do warming and precipitation change alter coflowering network structure?* We predicted that warming and drying would shorten the flowering season. This compression of the season would increase the amount and diversity of coflowering (network strength and weighted degree) and the number and proportion of coflowering species pairs (network size and connectance), while reducing the extent to which flowering communities are divided into distinct groups (modularity).
2. *How do phenological functional groups, growth forms, and origins (native vs. nonnative) drive climate-induced changes in coflowering network structure?* We predicted that hotter and drier conditions would primarily alter network structure through differential responses between phenological functional groups, whereby species belonging to the early-season functional group would not change their flowering phenology while those in the late-season functional group would shift flowering earlier, increasing overlap between early- and late-season species. We expected growth form and species origin to have comparatively limited effects, because while species within these groups may differ in their phenological sensitivities, their flowering phenologies are scattered across the growing season.
3. *How do climate conditions influence the flowering phenology and seed production of plants in early- and late-season phenological functional groups?* We predicted that flowering onset of early-season species would be insensitive to climate due to strong photoperiodic constraints, but expected flowering duration to extend under favorable cool– wet conditions. In contrast, we predicted that late-season species would be sensitive to climate throughout their flowering season, with warmer and drier conditions advancing flowering onset, peak, and end, because declining soil moisture would cause stress-induced flowering. For plants of both functional groups, we predicted that longer flowering durations that start earlier and end later would increase seed production.
4. *How do species vary in their phenological responses to warming and precipitation change?* Because species responses are idiosyncratic, we addressed this question without *a priori* hypotheses.

## Methods

We examined the effects of warming and altered precipitation on flowering phenology in experimental communities of native and non-native annuals from California serpentine grassland. Our study took place in 216 outdoor mesocosms situated within a ∼0.5-ha field at the Carnegie Institution for Science in Stanford, California, USA (37.43°N, 122.18°W).

### Study system

Most of California’s grasslands are dominated by non-native grasses, and have limited pollinator diversity. However, California’s widely dispersed patches of low-nutrient serpentine soils support diverse native forb and pollinator communities (Huenneke et al. 1990; Brennan 2022; Nelson et al. 2022). These (typically shallow) soils favor annual plant species, most of which can be broadly grouped into two functional groups that partition resources temporally: early-season annuals and late-season annuals. Both early- and late-season annuals germinate with the first autumn rains, but early-season annuals flower in spring and senesce before summer drought, whereas late-season annuals develop deep taproots throughout the spring and rely on deep soil moisture to survive the predictable summer drought (Pitt and Heady 1978; Gulmon et al. 1983; Jackson and Roy 1986; Mooney et al. 1986; Hooper and Vitousek 1998; Harrison and Viers 2007; Levine et al. 2022). As these groups use resources at different times, competition is strongest within rather than between groups, and this temporal partitioning promotes coexistence (Hooper and Vitousek 1998; Levine & HilleRisLambers 2009; Hooper and Dukes 2010; Levine et al. 2022). The region’s Mediterranean climate features cool, wet winters and hot, dry summers.

Although changes to mean temperature and precipitation are projected to be modest in the coming decades (Neelin et al. 2013; Diffenbaugh et al. 2015; Simpson et al. 2016), interannual precipitation variability is high and expected to increase, yielding more frequent extremely wet or dry years (Neelin et al. 2013; Yoon et al. 2015; Swain et al. 2018). Historically, droughts have been most likely to occur when low precipitation co-occurs with warm conditions, and years that are both warm and dry are increasingly common across the state (AghaKouchak et al. 2014; Diffenbaugh et al. 2015).

### Experimental set-up

We seeded communities of native serpentine annual plants and common non-natives into 0.018 m² mesocosms, and exposed them to a broad range of climate treatments, spanning ambient and warmed temperature treatments and ambient precipitation, precipitation removal, and precipitation addition treatments (*Fig. S1*).

We initially included 14 species in the experiment (*Table 1*). We selected widespread and locally abundant species with seeds that were practical to collect in a field environment and prioritized the inclusion of natives and non-natives of different functional groups and growth forms. All seeds were collected from the nearby Jasper Ridge Biological Preserve (’Ootchamin ’Ooyakma; 37.40°N, 122.24°W). As this experiment was originally designed to test how functional similarity among competitors and climate conditions shape native–invader coexistence, we grew these seeds in communities that included both a native and a non-native component. The native component of the community included either all of the early-season native annual species, all of the late-season native annual species, a combination of all early- and late-season native species, or a control without any native species. The non-native component of the community included either all of the early-season non-native annual species, all of the late-season non-native annual species, or a control without any non-native species. While this planting strategy theoretically yielded communities of 3–11 species in each mesocosm, variable germination and survival rates meant that some communities had as few as one species that survived to flower. Furthermore, as only 14 *Erodium botrys* and no *Dittrichia graveolens* survived to a reproductive life stage, these species were excluded from the analysis (*Table 1*).

**Table 1.**
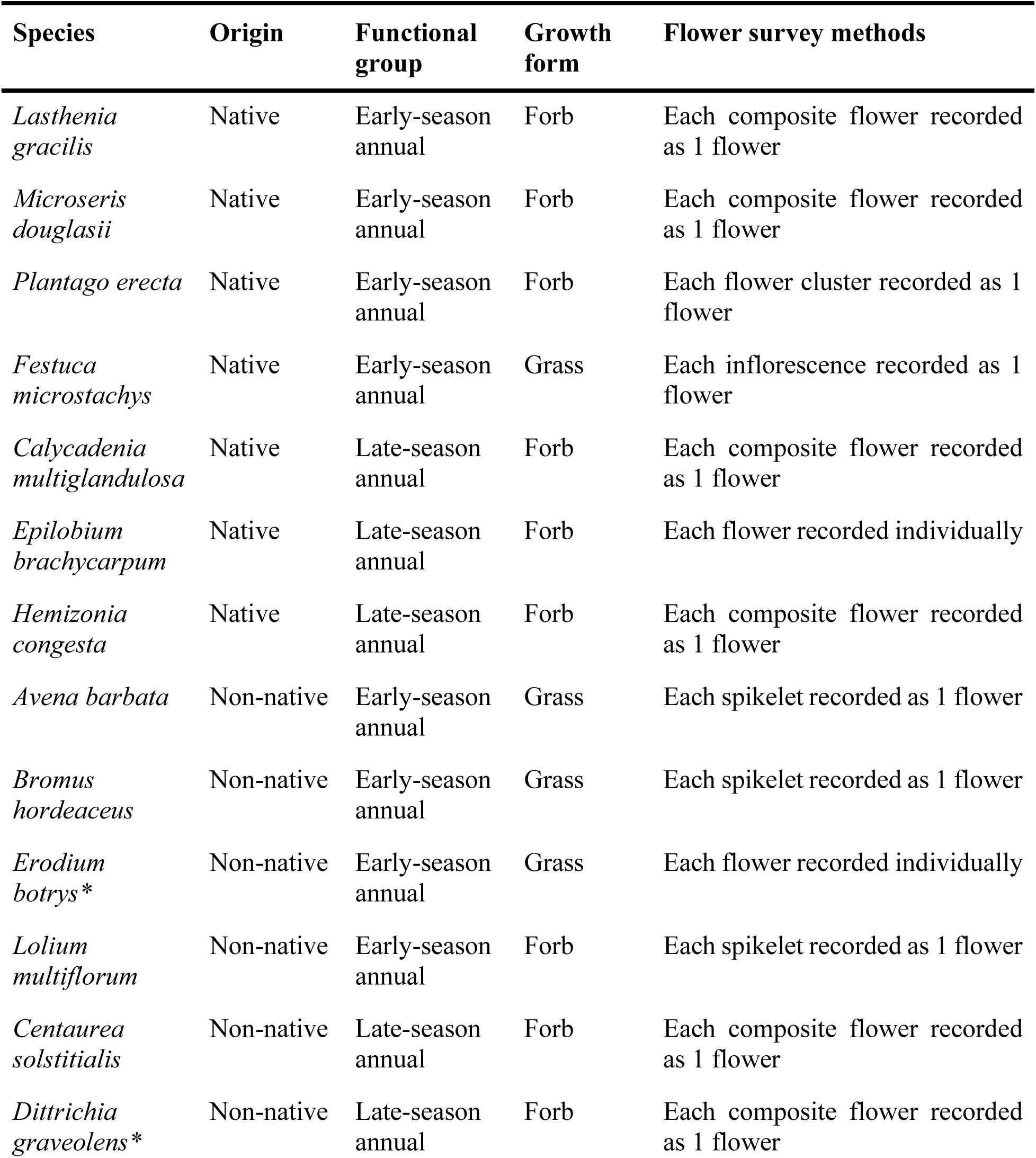
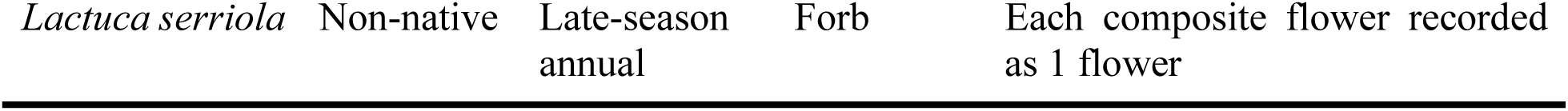
Species classified by origin, phenological functional group, and growth form. All forbs in this study are insect pollinated and all grasses in this study are wind-pollinated. Species marked with an asterisk were excluded from the analysis due to low germination and survival rates.

We constructed the mesocosms using 15-cm-diameter, 100-cm-tall PVC pots filled with serpentine soil collected from a grassland in Morgan Hill, California, USA (37.18°N, 121.68°W), managed by the Kirby Canyon Recycling & Disposal Facility. On 20–21 November 2023, we seeded the pots with experimental communities at densities approximating natural serpentine assemblages (Gulmon et al. 1983). We then allowed them to develop without further manipulation of species composition or density, except for removing any species not included in the experimental design. We organized the mesocosms into 18 blocks of 12 mesocosms, yielding 216 mesocosm communities, and randomized both the climate treatments assigned to each block and the community composition assigned to each mesocosm within a block.

We subjected the mesocosm communities to a range of simulated climate conditions. We manipulated temperature by surrounding some blocks with open-top enclosures made of reflective bubble wrap or transparent plastic, which increased ambient temperatures by ∼1.5°C and ∼6°C, respectively (*Fig. S2*). To create lower-moisture treatments, we built rainout shelters constructed from frames fitted with U-shaped transparent plastic troughs that intercepted 30% or 60% of ambient rainfall and deposited it into collection buckets. To create higher-moisture treatments, we supplemented rainfall at the end of the rainy season by hand-watering the mesocosms with the collected rainwater twice weekly for three to nine weeks, applying an additional 30%, 60%, or 90% of ambient rainfall. Together, these structures produced three temperature treatments (ambient, warm, hot) and precipitation treatments ranging from -60% to +90% of the ambient rainfall received by the mesocosms throughout the year (*Fig. S2*). We measured soil surface temperature and volumetric water content (VWC) from 19 March–20 June 2024 in all no-plant control mesocosms and all single functional–group mesocosms (i.e., those containing only early-season natives, early-season non-natives, late-season natives, or late-season non-natives) in the - 60%, control, and +90% precipitation treatments (54 mesocosms total) with TMS-4 dataloggers (TOMST, Prague, Czech Republic). We then used a random forest model to estimate the average climate conditions experienced by the remaining 162 mesocosms over the same period (see Supplemental Information for details).

### Flowering phenology and seed production

We surveyed flowering phenology by counting the number of flowers of each species in each mesocosm every 3–10 days from the day when flowering was first observed in the experiment on 23 February 2024 to the end of the experiment on 29 September 2024 (*Fig. 1*). For species for which recording individual flowers would have been impractical, we recorded each spikelet, inflorescence, or composite flower as a single flower (*Table 1*). To estimate seed production for each species in each mesocosm, we collected mature seeds daily throughout the spring and summer and measured their total mass. We then weighed and counted a subset of the collected seeds to calculate the average mass per seed and used the average mass per seed and the total seed mass of each species in each mesocosm to estimate the total number of seeds produced.

**Figure 1.**
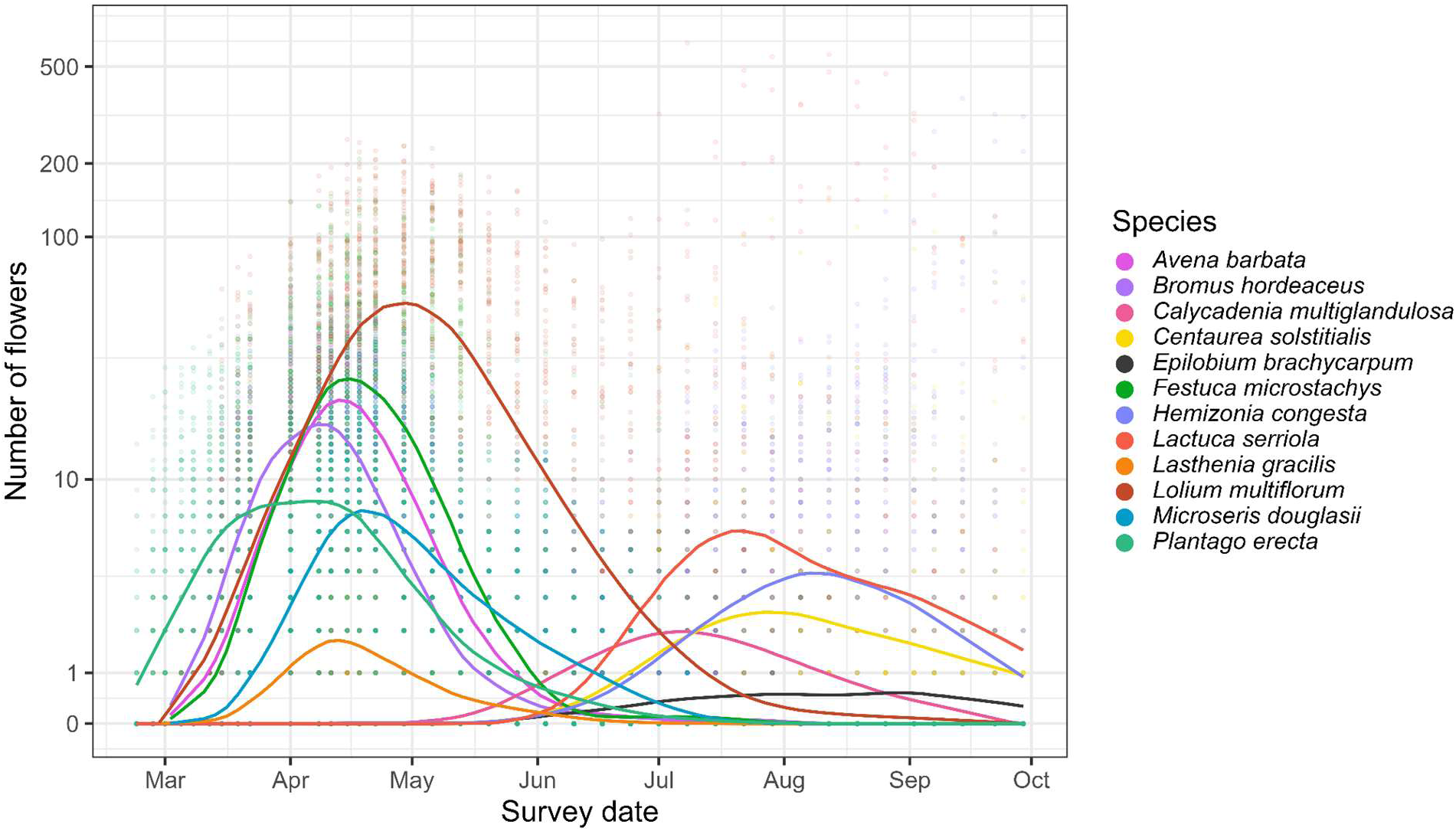
The number of flowers of each species throughout the February 23–September 29 2024 survey period. Shown are points representing the number of flowers recorded for each species in a given mesocosm on each survey date and smoothed lines (LOESS fits, degree = 1, span = 0.25) highlighting temporal trends in flowering across all climate treatments.

### Coflowering network analysis

We investigated how climate conditions influence coflowering patterns among California serpentine annuals by constructing coflowering networks based on Schoener’s index of niche overlap (Schoener 1970; Arceo-Gómez et al. 2018). When used in this way, Schoener’s index (SI) quantifies the degree of overlap in flowering phenology between pairs of species, incorporating both the intensity (number of flowers produced) and frequency (number of survey days in which flowers were observed) of flowering, meaning that higher SI values indicate species pairs that flower simultaneously for longer periods and with greater intensity (i.e., species whose curves have greater overlap in *Fig. 1*).

To examine the effects of climate on coflowering network structure, we first assigned the mesocosms to four climate groups based on natural break points from a visual analysis of springtime surface temperature and soil volumetric water content (hereafter, “soil moisture”): cool–dry, cool–wet, warm–dry, and warm–wet (*Fig. S5*). We then bootstrapped flower abundances in these treatments by randomly selecting groups of mesocosms within each climate treatment and pooling their species-level flower counts on each day throughout the survey period and calculating the SI for all species pairs and derived mean SI values and 95% confidence intervals (*Figs. S6- S8*). We repeated this procedure 1000 times, then used the average SI values to construct coflowering networks for each community using the R package *igraph* (Csárdi et al. 2025).

We characterized network structure by calculating average coflowering strength (the intensity and duration of coflowering between species pairs), average weighted degree (the strength-weighted number of coflowering interactions with a given species, which provides a measure of the diversity of coflowering interactions the species experiences), modularity (the degree to which groups of species interact more strongly within the group than with other species in the network), network size (number of coflowering species pairs), and network connectance (the proportion of realized coflowering pairs relative to the theoretical maximum).

To test how warming, precipitation change, and species attributes altered the average strength of coflowering between pairs of species, we used linear mixed-effects models with SI as the response variable. Predictor variables included climate treatment (cool–dry, cool–wet, warm– dry, or warm–wet), functional group combination (early–early, late–late, or early–late), growth form combination (grass–grass, forb–forb, or grass–forb), and origin combination (native–native, non-native–non-native, or native–non-native). The species pair was included as a random effect to account for repeated measures across the networks. To evaluate how climate and species attributes affected the diversity of coflowering relationships experienced by individual species, we calculated the weighted degree of each species in the network and modeled it as the response variable in linear mixed-effects models. Predictor variables included the climate treatment and species’ functional group, growth form, and origin. Species identity was included as a random effect.

We assessed homogeneity of variances and normality of residuals using residual-versus-fitted plots and normal Q–Q plots. After verifying assumptions, we evaluated model fit using piecewise structural equation modeling with R package *piecewiseSEM* (Lefcheck et al. 2016), which allowed us to quantify the variance explained by fixed effects alone versus the full model including random effects. Fixed effects were then analyzed with ANOVA, and pairwise comparisons among climate, growth form, and origin treatments were conducted using the R package *emmeans* (Searle et al. 2020). For significant main effects and interactions, we compared estimated marginal means using Tukey-adjusted p-values and Kenward–Roger degrees of freedom.

### Climate effects on flowering phenology and seed production

To examine how warming and precipitation change altered the onset, peak, end, and duration of flowering and the consequences of both climate and flowering phenology on seed production, we applied piecewise structural equation modeling using the R package *piecewiseSEM* (Lefcheck et al. 2016). These models allowed us to evaluate the effects of the average springtime soil moisture and temperature experienced by a given mesocosm on seed production both directly and through their indirect effects mediated by flowering phenology.

For a given species in a given mesocosm, we quantified the stages of flowering phenology by calculating the day of year marking the onset of flowering, peak of flowering, and end of flowering, as well as the total flowering duration. Onset and end of flowering were determined from daily flower counts as follows: the onset was the first day on which one or more flowers were observed, provided it was followed by at least one additional day with flowers; the end was the last day on which flowers were observed, provided it was preceded by at least one other day with flowers; peak flowering was estimated by fitting a LOESS smoothing curve (span = 0.3) to daily flower counts for each species-mesocosm combination and identifying the day with the highest predicted flower number. In cases where only a single flowering observation occurred, the onset, peak, and end of flowering were all recorded on that same day, and the flowering duration was set to one day. Flowering duration was calculated as the interval between the onset and end of flowering. We additionally included both the total end-of-season biomass and the maximum number of flowers observed for each species in each mesocosm in the model to account for the potential influence of climate on these plant size variables and for potential allometric influences on phenology and seed production. Although we initially included both block and mesocosm as additional random effects, they explained minimal variance and caused convergence issues, so we removed them from the final models.

Given the strong flowering modularity within early- and late-season functional groups and the distinct responses of early- and late-season species to different climate treatments in our analysis of coflowering networks, we developed separate SEMs for early-season and late-season annuals. This allowed us to capture functional group–specific effects of springtime temperature and soil moisture on flowering phenology and reproductive performance. For each model, we assessed global model fit using Fisher’s C test and included species identity as a random effect.

### Species-specific variation in flowering phenology responses to warming and precipitation change

To examine interspecific variation in warming- and precipitation-induced changes in flowering phenology, maximum flower count, and seed production, we ran a series of linear mixed-effects models (LMMs) with R package *afex* (Singmann et al. 2016). For each, we fit the onset, peak, end, or duration of flowering, the maximum flower count, or the number of seeds produced as the outcome variable and the species and climate treatment as the fixed effect variables, with block and mesocosm number as random variables. We assessed whether the models met assumptions, evaluated model fit, and analyzed our results as above, except that we log-transformed maximum flower number and seed production to meet the model assumptions. All statistical analyses and visualizations were conducted in R (RStudio v.2023.03.0+386, R Core Team 2023), and the full datasets and R scripts are available on Figshare (Nebhut and Dukes 2026).

## Results

Community-wide flowering displayed pronounced temporal structuring. The total number of flowers across all species followed a bimodal pattern driven by phenological differences between functional groups, with early-season species flowering mainly in spring and late-season species flowering primarily in summer (*Fig. 1*). As a result, the coflowering networks of each climate treatment contained two distinct coflowering modules: one composed of early-season annuals and one of late-season annuals (*Fig. 2*).

**Figure 2.**
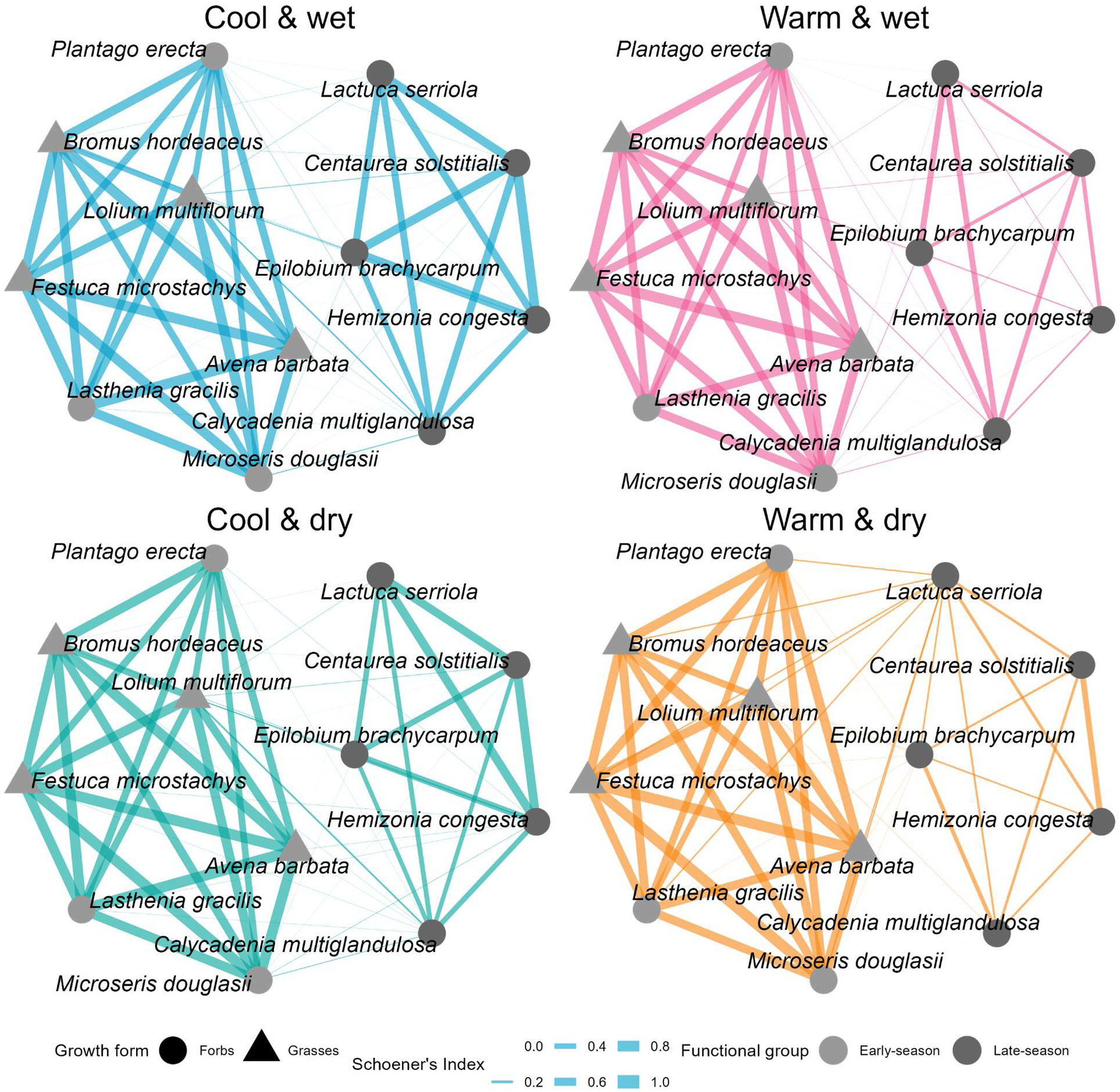
Coflowering networks of serpentine grassland annual communities in the four climate treatments (teal: cool–dry, blue: cool–wet, orange: warm–dry, and pink: warm–wet). Each node (point) represents a plant species, and the links (lines) between them represent coflowering interactions, with the thickness of the lines indicating the amount of flowering overlap for a given pair of species (Schoener’s index). Early-season annuals are represented in light grey and late-season annuals in dark grey; forbs are represented as circles and grasses as triangles.

### Warm conditions altered coflowering network structure by diminishing coflowering among late-season annuals

Warming reduced the average strength and diversity of coflowering interactions as well as network-wide modularity, network size, and connectivity (*Table 2*). These results indicate that California serpentine grassland communities exhibit less coflowering and less distinct functional groups in warm conditions than in cool ones.

**Table 2.** Network metric values for coflowering networks in the four climate treatments. Network metric Climate treatment

| Network metric | Climate treatment |  |  |  |
| --- | --- | --- | --- | --- |
|  | Cool & dry | Cool & wet | Warm & dry | Warm & wet |
| Average strength | 0.29 | 0.3 | 0.25 | 0.25 |
| Average weighted degree | 3.21 | 3.27 | 2.8 | 2.77 |
| Network modularity | 0.36 | 0.39 | 0.2 | 0.27 |
| Network size | 64 | 65 | 60 | 61 |
| Network connectance | 0.97 | 0.98 | 0.91 | 0.92 |

While early-season annuals experienced consistent coflowering relationships across climate treatments, warming weakened the coflowering strength and diversity experienced by late-season annuals. Coflowering strength (SI) was highest between pairs of early-season species, reflecting substantial overlap in their flowering phenologies, and lowest between species belonging to different functional groups, which exhibited minimal temporal overlap in flowering across all climate treatments. Late-season pairs showed intermediate SI values that were sensitive to climate treatment: coflowering strength was highest in the cool–wet condition and lowest in both warm treatments (*Fig. 3a*). The diversity of species’ coflowering interactions (weighted degree) showed similar functional group–specific patterns. Early-season annuals had consistently high coflowering diversity, while late-season species exhibited reduced coflowering diversity under warm conditions (*Fig. 3b*). In contrast, origin (native or nonnative) and growth form (grass or forb) had no effect on the average coflowering strength of species pairs or coflowering diversity experienced by individual species (*Table 3*).

**Figure 3.**
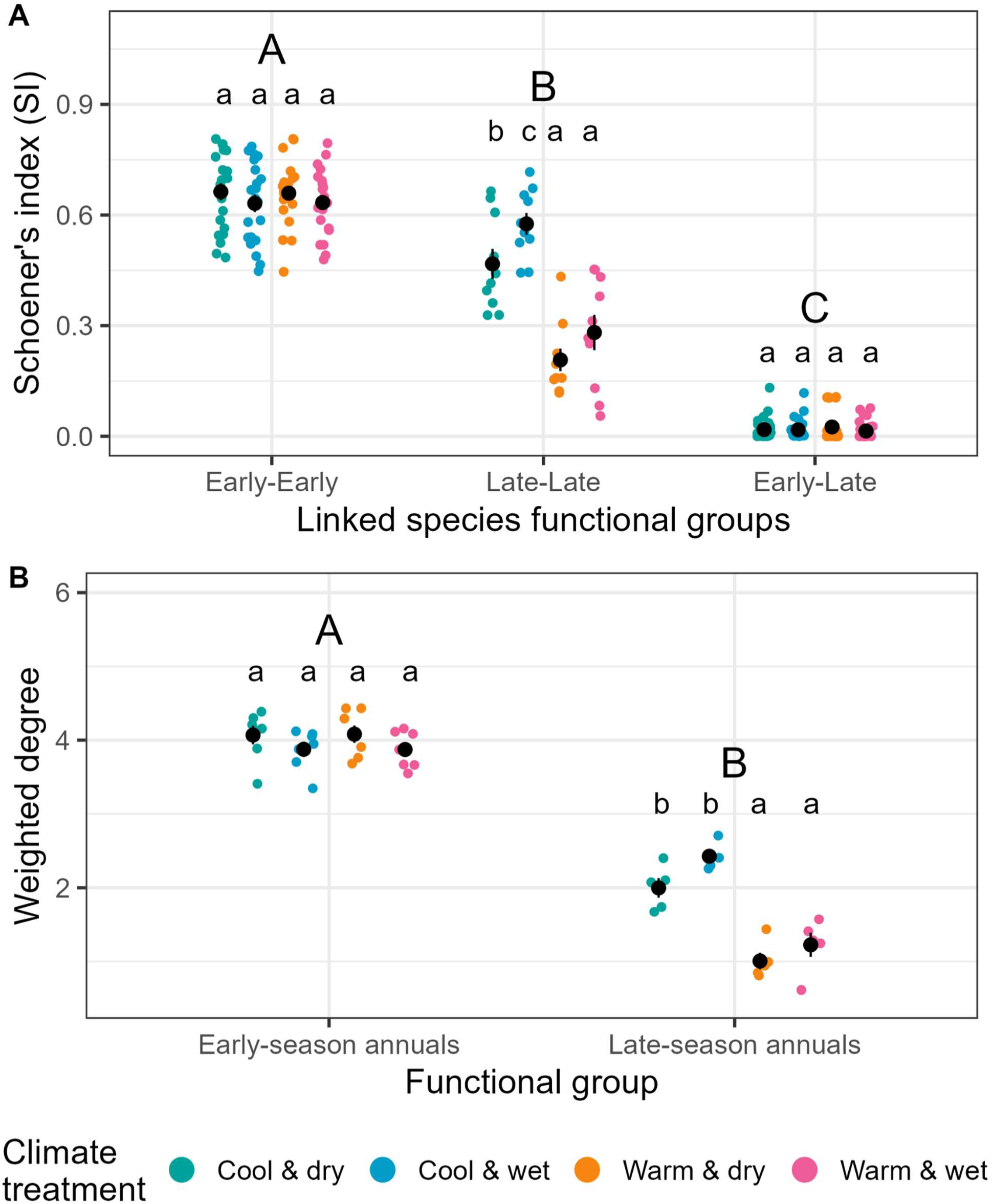
A) Differences in the strength of coflowering for a pair of species (Schoener’s index; SI) and B) the diversity of coflowering interactions of each species (weighted degree) in the four climate treatments (cool & dry, cool & wet, warm & dry, and warm & wet). Shown are the data, the means ± SE, and post-hoc test results; significant differences between functional group combinations or functional groups are indicated in different uppercase letters and significant differences in climate treatments within functional group combinations or functional groups are indicated in different lowercase letters.

**Table 3.**
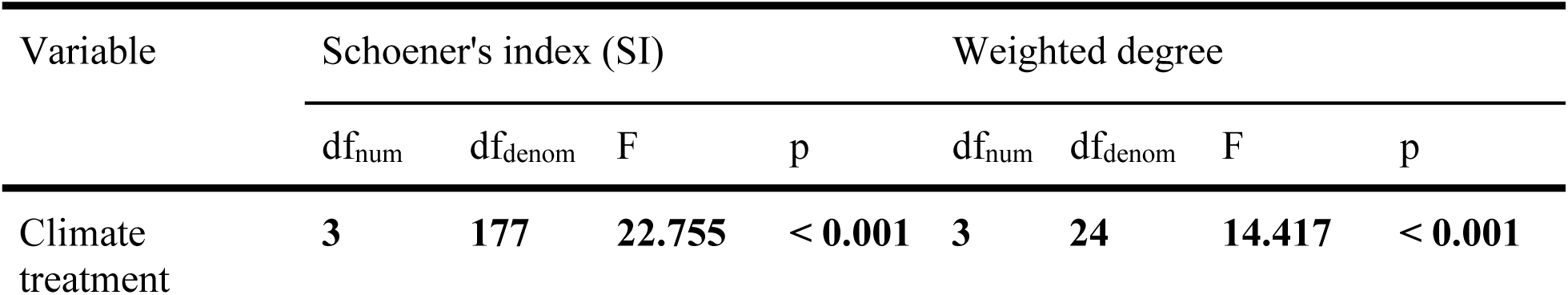
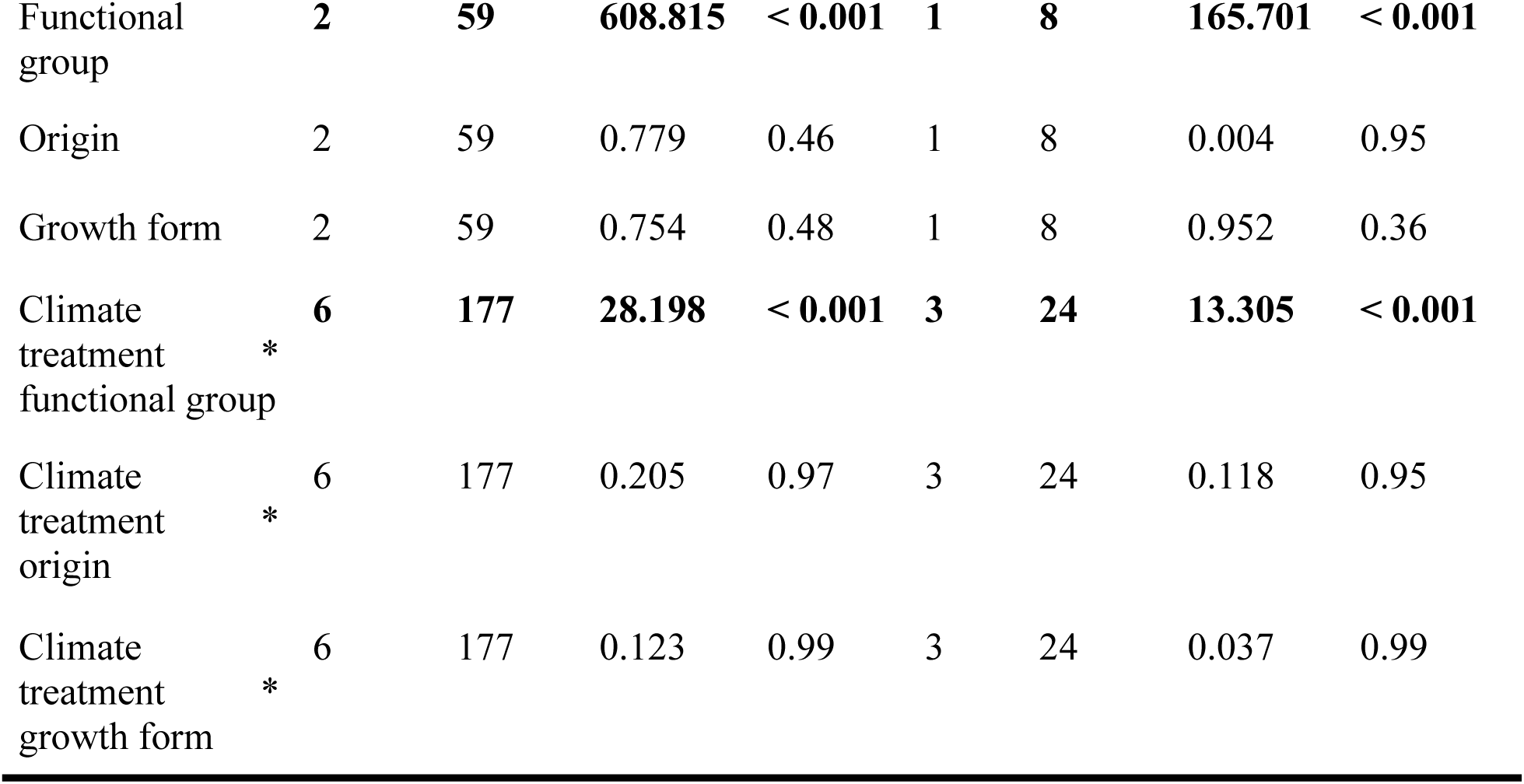
Statistical results from linear mixed-effects models examining how climate treatment, functional group, origin, growth form, and their interactions influenced coflowering strength (SI) and diversity (weighted degree). Significant p-values are shown in bold. The fixed effects in the model for coflowering strength and diversity explained 93.7% and 94.1% of the variance (marginal R²), respectively, with both the random and fixed effects together accounting for 96.4% and 96.1% of the variance (conditional R²), respectively.

| Variable | Schoener's index (SI) |  |  |  | Weighted degree |  |  |  |
| --- | --- | --- | --- | --- | --- | --- | --- | --- |
|  | df<sub>num</sub> | df<sub>denom</sub> | F | p | df<sub>num</sub> | df<sub>denom</sub> | F | p |
| Climate treatment | **3** | **177** | **22.755** | **< 0.001** | **3** | **24** | **14.417** | **< 0.001** |
| Functional group | 2 | 59 | 608.815 | < 0.001 | 1 | 8 | 165.701 | < 0.001 |
| Origin | 2 | 59 | 0.779 | 0.46 | 1 | 8 | 0.004 | 0.95 |
| Growth form | 2 | 59 | 0.754 | 0.48 | 1 | 8 | 0.952 | 0.36 |
| Climate treatment * functional group | 6 | 177 | 28.198 | < 0.001 | 3 | 24 | 13.305 | < 0.001 |
| Climate treatment * origin | 6 | 177 | 0.205 | 0.97 | 3 | 24 | 0.118 | 0.95 |
| Climate treatment * growth form | 6 | 177 | 0.123 | 0.99 | 3 | 24 | 0.037 | 0.99 |

### Climate conditions altered seed production indirectly via functional group–dependent changes to flowering phenology

Early- and late-season species likewise differed in how average springtime temperature and soil moisture, plant size (vegetative biomass and maximum flower count), and flowering phenology interacted, and in what consequences these variables had on seed production (*Fig. 4*; *Table S1*). For early-season annuals, average springtime temperature and soil moisture influenced seed production both directly and indirectly through their effects on the timing of the peak and end of flowering (Fisher’s C = 0.834; df = 2; p = 0.66). End-of-season vegetative biomass and the maximum number of flowers produced by early-season annuals were not directly affected by average springtime temperature or soil moisture, although early-season annuals with greater vegetative biomass produced a higher maximum number of flowers. Onset of flowering was similarly unaffected by climate but occurred earlier in early-season annuals with more biomass and more flowers, while peak flowering occurred earlier in early-season annuals that began flowering earlier, in those with more flowers, and in warmer, wetter, or both warmer and wetter conditions. End of flowering occurred earlier in early-season annuals that reached peak flowering earlier, but later in early-season annuals with more biomass, in those producing a greater maximum number of flowers, and in wetter conditions. Flowering duration increased when early-season annuals both began earlier and ended later, and was longer in plants with greater biomass or a smaller maximum number of flowers. Early-season annual seed production was driven both directly by climate and indirectly by phenology: early-season annuals that began flowering earlier or ended later produced more seeds. After accounting for these effects, shorter flowering durations were also associated with higher seed production. Early-season annuals with greater vegetative biomass and those that produced a higher maximum flower count had higher seed output, and early-season annuals in drier conditions or in conditions that were both warmer and wetter produced more seeds.

**Figure 4.**
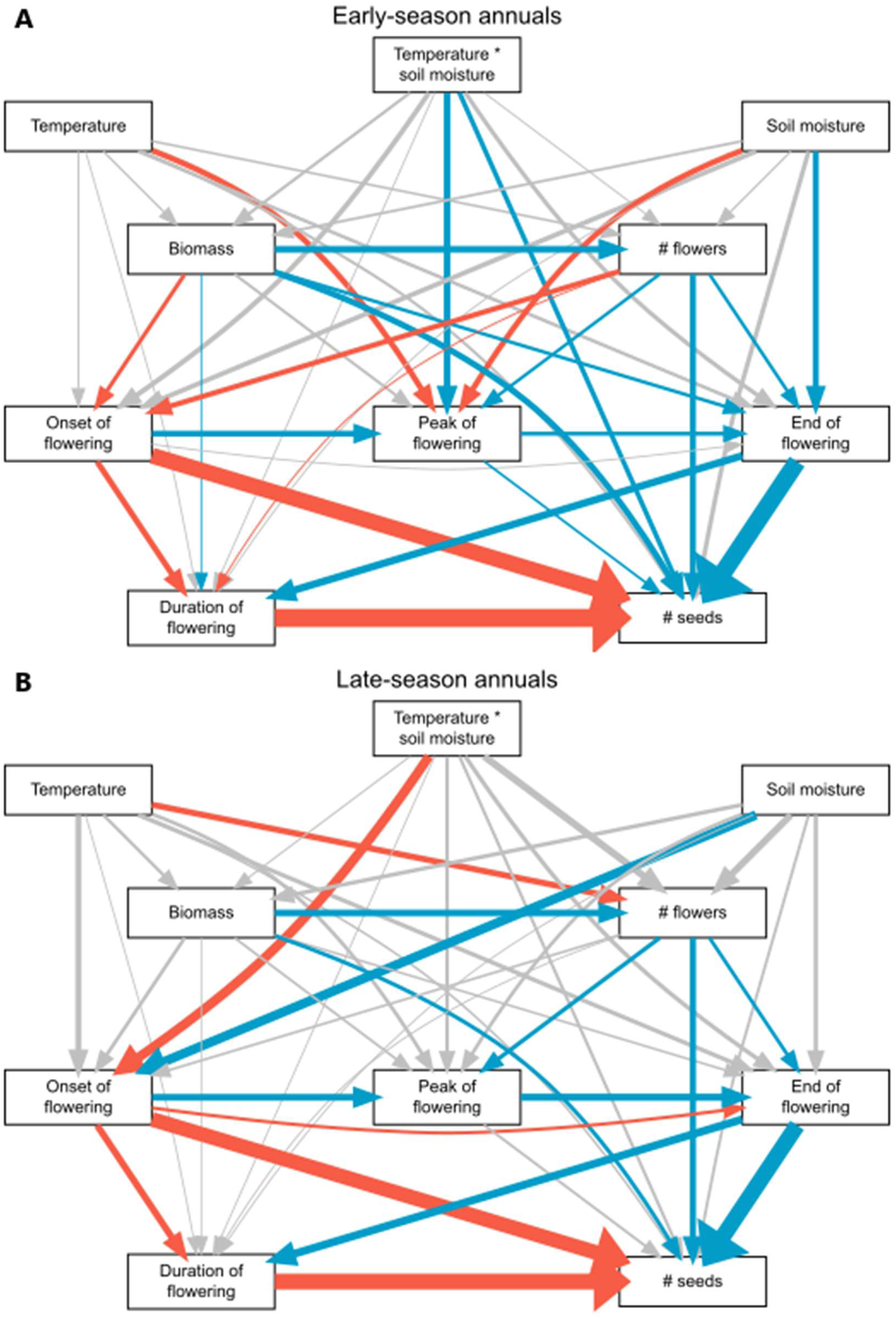
Structural equation model linking climate conditions to plant size variables, flowering phenology, and seed production for A) early-season annuals and B) late-season annuals. Single-headed arrows describe directional dependence relationships between these variables, and the thickness of the lines indicate the standardized path coefficients. Blue arrows indicate a positive relationship (an increase in the explanatory variable results in an increase in the response variable), red arrows indicate a negative relationship (an increase in the explanatory variable results in a decrease in the response variable), and grey arrows indicate a non-significant relationship.

For late-season annuals, climate influenced seed production only indirectly via its effects on flowering phenology and flower production (Fisher’s C = 2.813; df = 2; p = 0.25). Average springtime temperature and soil moisture did not affect vegetative biomass production, but late-season annuals with greater biomass produced greater maximum numbers of flowers, and those in cooler conditions also produced more flowers. The onset of late-season annual flowering occurred earlier in wetter or both cooler and wetter conditions, and late-season annuals that began flowering earlier or produced more flowers reached peak flowering earlier, while the end of flowering occurred earlier in late-season annuals that began flowering later, reached peak flowering later, or produced a greater maximum number of flowers. Flowering duration increased when late-season annuals both started earlier and ended later, but was otherwise unaffected by climate or plant size variables. Similar to the early-season annuals, late-season annual seed production was strongly influenced by the onset, end, and duration of flowering. Late-season annuals that began flowering earlier or ended later produced more seeds, but after accounting for these effects, shorter flowering durations were associated with greater seed output. Late-season annuals with more vegetative biomass and those with more flowers also produced more seeds. Unlike our findings for early-season annuals, however, we found that average springtime temperature and soil moisture had no direct effect on the seed production of late-season annuals.

### Species differ in their flowering phenology and climate responses

Species differed in their flowering phenology, maximum flower count, seed production, and responses to climate treatments (*Fig. 5*; *Tables S2–S3*). On average, *Plantago erecta* initiated flowering earliest (5 March), whereas *Lactuca serriola* flowered latest (5 July). *Bromus hordeaceus* completed flowering first (8 May), and *Centaurea solstitialis* the latest (18 August). Flowering duration ranged from an average of 13.5 ± 5.8 days in *Epilobium brachycarpum* to 84.2 ± 2.5 days in *Lolium multiflorum*. Reproductive output also differed substantially, with *Lasthenia gracilis* producing the lowest maximum number of flowers (2.9 ± 1.1 flowers on average) and *Lolium multiflorum* the greatest (103.9 ± 1.1 flowers), and with *Calycadenia multiglandulosa* producing the fewest seeds (8.5 ± 1.4 seeds) and *Lolium multiflorum* the most (461.3 ± 1.1 seeds). The species also varied in their phenological and reproductive responses to the climate treatments (*Fig. 5*; *Tables S2–S3*). For example, *Microseris douglasii* produced similar numbers of flowers and seeds across treatments; however, onset, peak, and end of flowering were progressively earlier in warm–dry, warm–wet, and cool conditions, and flowering duration was longest in cool–wet and shortest in warm–dry conditions. In contrast, *Hemizonia congesta* initiated flowering later in cool–wet than warm–wet conditions, reached peak and ended flowering later in cool than warm climates, and had longer flowering durations, greater flower production, and higher seed production in cool–wet than in warm–dry conditions.

**Figure 5.**
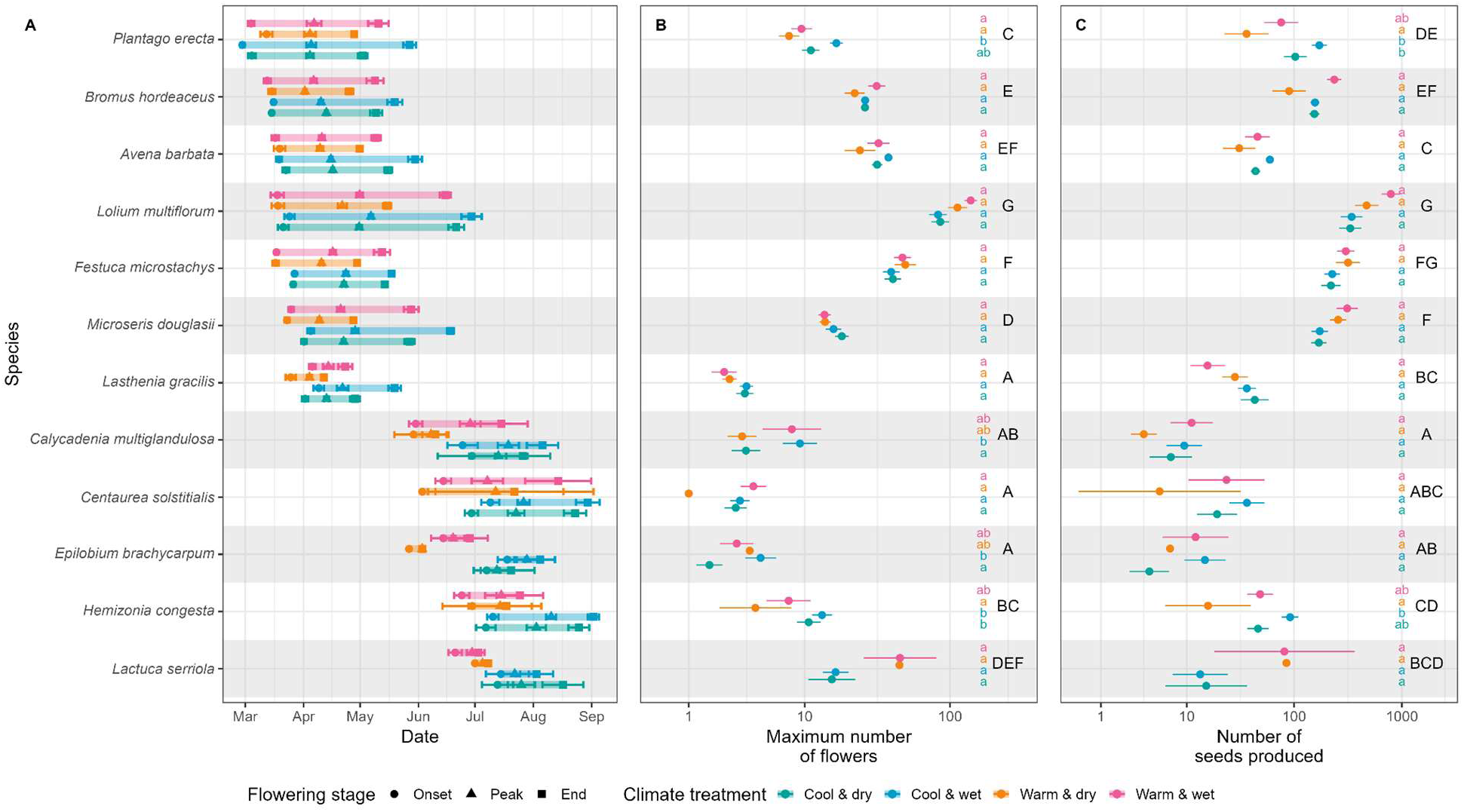
Flowering phenology by species, including A) the average (mean ± SE) onset, peak, and end of flowering, B) the maximum number of flowers of each species in each mesocosm observed on a single sampling day (mean ± SE) and C) the total number of seeds produced by each species in each mesocosm throughout the year (mean ± SE). For panel A, significant differences between species and treatments within species are in Table S2, and for panels B and C, significant differences between species are indicated by different uppercase letters and significant differences in climate treatments within species are indicated by different lowercase letters.

## Discussion

California serpentine grassland communities exposed to warmer springtime average temperatures showed reduced and less modular coflowering, driven largely by weakened and less diverse coflowering among late-season annuals. Springtime average soil moisture altered these patterns: late-season plants coflowered weakly with each other under both warm conditions regardless of soil moisture, but more strongly when conditions were wet and cool than when they were dry and cool. Growth form and origin did not influence species’ average coflowering strength or diversity, or how sensitive these metrics were to changes in climate. Flowering phenology affected seed production in both early- and late-season functional groups. However, climate effects on flowering phenology differed between these groups. In early-season plants, flowering onset was insensitive to climate, but cooler temperatures or wetter soil delayed peak flowering and flowering end. In contrast, late-season plants advanced flowering onset under warmer and drier conditions, while climate-driven changes in peak flowering and flowering end occurred only indirectly through temperature-driven changes in maximum flower number. Despite these patterns in functional group responses to altered climate conditions, substantial species-level variation in phenological responses ultimately shaped how warming and precipitation changes restructured coflowering networks and the seed production of individual species.

### Warming-driven changes in coflowering network structure are driven by reductions in coflowering within the late-season functional group

We anticipated that warming and drying would cause species to increasingly overlap in their flowering phenology, thereby increasing the strength and diversity of coflowering and the number and proportion of coflowering species pairs. Instead, we found the opposite: under warm conditions, and often under conditions that were both warm and dry, coflowering strength and diversity were reduced, paired with decreases in network size and connectivity. Network modularity was likewise reduced by warm and dry conditions.

This pattern of weakened phenological overlap and reduced modularity under warm conditions was caused by weakened and less diverse coflowering among late-season annuals. These findings parallel those of Fisogni et al. (2022), who observed reduced coflowering at warmer sites along an elevational gradient and Park et al. (2025), who found that reduced late-season coflowering is characteristic of warming plant communities across California. This suggests that the divergence in coflowering communities in response to climate change may be widespread, though increasing overlap can also occur (Diez et al. 2012).

Notably, although we expected warming and drying to increase community-wide phenological overlap by promoting greater coflowering between early- and late-season species, neither treatment, alone or combined, altered coflowering strength between early- and late-season species pairs. This means that in these California serpentine grassland annuals, pre-existing phenological separation between early- and late-season functional groups (Chiariello 1989) and distinct climate responses (e.g., warming and drying advancing the end of flowering in early-season species but the onset of flowering in late-season species) maintained the already-minimal coflowering between groups. Thus, the observed loss of modularity in warmed communities reflects the erosion of coflowering strength within the late-season coflowering module, not increased synchrony between functional groups, as originally hypothesized. This suggests that early-season plants may maintain their current coflowering relationships under climate change, while late-season species may experience reduced coflowering.

Growth form and origin (native vs. non-native species) did not influence coflowering strength, coflowering interaction diversity, or their responsiveness to altered climate conditions. This contrasts with studies showing greater phenological responsiveness of forbs relative to grasses (Iverson et al. 2009) and of non-native species relative to natives (Willis et al. 2010; Lesica and Kittelson 2010; Calinger et al. 2013; Wolkovich et al. 2013; Zettlemoyer et al. 2019). However, these studies examined direct phenological responses rather than their consequences for coflowering structure. Whether changes in flowering phenology alter patterns of coflowering depends on species having asynchronous responses to changing conditions (Fisogni et al. 2022). That is, coflowering relationships can change only when species respond differently to altered conditions – either by shifting flowering phenology in different directions or by different magnitudes – and when species are sufficiently clustered within the flowering season such that increases in coflowering with some species are not offset by decreases with others. In this case, although climate sensitivity may have differed among species of varying growth forms and origins, their flowering periods were sufficiently dispersed across spring and summer that any phenological shifts were insufficient to alter the average strength or diversity of coflowering experienced within these groups.

### Early-season species extend flowering in favorable conditions, while late-season species flower under stress

Species belonging to the early- and late-season phenological functional groups differed in their phenological responses to average springtime temperature and soil moisture. Early-season species exhibited climate-insensitive flowering onset, consistent with photoperiodic control (Hirose et al. 2005; Jackson 2008). Such species typically show weak phenological responses to changing climate conditions and are thought to be disadvantaged relative to more responsive competitors, as they may miss opportunities to extend their growing season with earlier flowering (Zeng et al. 2025). Late-season species showed strong phenological sensitivity to changing climate conditions in their flowering onset, which was advanced by drying and by combined warm–dry conditions, with their peak and end of flowering advancing in parallel with flowering onset. Cooler temperatures prolonged flowering indirectly by enabling greater floral production before senescence, but otherwise, wetter and cooler conditions did not enable late-season annuals to delay their flowering peak or end. These patterns are consistent with stress-induced flowering, in which plants accelerate reproduction when resource stress prevents continued growth (Wada and Takeno 2010; Riboni et al. 2014; Takeno 2016).

These results contrast with other temperate-system studies in which early-flowering species responded more strongly to warming than their later-flowering counterparts (Price and Waser 1998; Fitter and Fitter 2002; Miller-Rushing and Primack 2008; Moore and Lauenroth 2017). In most temperate systems, early-flowering plants stand to benefit more from earlier growth and flowering than late-flowering species, as advancing their phenology allows them to capitalize on higher light availability before canopy closure (Calinger et al. 2013). In contrast, light is rarely limiting in low-statured California serpentine grasslands (Turitzin et al. 1978), so early advancement likely confers fewer benefits. Likewise, wintertime growth is constrained by short days and low irradiance, restricting the potential for earlier onset even under warming (Chiariello 1981, 1989). These limitations lift as the season progresses, and early-season annuals tend to reproduce for longer during favorable years, enabling greater total annual carbon and nitrogen uptake (Woodmansee and Duncan 1980). Conversely, late-season annuals may benefit from delaying the shift from vegetative growth to reproduction for as long as conditions allow. Therefore, late-season plants in cool-wet environments can sustain growth for longer into the summer and delay the onset of flowering relative to late-season plants in hot-dry conditions. In either case, flowering begins once available soil moisture is depleted and subsequent flowering stages advance in parallel with flowering onset (Gulmon et al. 1983; Chiariello 1989).

Past studies in Mediterranean climates provide a mixed picture of how early-season species respond to climatic variation. Several investigations of early-season non-native grasses have reported little to no effect of either drying or increased water availability on flowering phenology (Gulmon 1979; Ewing and Menke 1983; Jackson and Roy 1986). In contrast, Petanidou et al. (1995), working in a comparable Mediterranean system, found that winter warming advanced flowering time, with the strongest effects observed in early-flowering taxa. Similarly, Van Dyke and Kraft (2025) documented substantial phenological shifts in response to reduced rainfall among multiple early-season serpentine annuals, including some species that did not show strong responses to drying in our experiment. Both methodological and environmental factors may help explain these discrepancies. Van Dyke and Kraft (2025) sampled flowering phenology more frequently (every 1–5 days) than the 3–10 day intervals used in this study, increasing the ability to detect subtle or early shifts in flowering (Miller-Rushing et al. 2008). As well, soil moisture remained relatively high across all of our experimental treatments during winter and early spring, and temperature differences between treatment groups were minimal during this wet, cloudy period because our passive heating chambers produced the strongest warming on sunny days (see Supplemental Information). As a result, climatic contrasts among treatments may have been too small to evoke phenological effects early in the season.

### Species-level responses shape coflowering networks

As we expected, we found substantial species-to-species variability in how warming and altered precipitation affected flowering phenology and seed production, including within functional groups. Indeed, while functional groups provided a useful framework for predicting general phenological responses to different climate conditions, substantial species-to-species variation within functional groups ultimately altered coflowering network structure by changing patterns of late-season coflowering (Aldridge et al. 2011; Diez et al. 2012; Fisogni et al. 2022). Early-season annuals exhibited relatively consistent and comparatively small phenological shifts across species, which helped maintain stable coflowering relationships under different climate regimes. In contrast, late-season annuals showed pronounced heterogeneity in their responses to temperature and soil moisture, leading to substantial climate-driven restructuring of their coflowering interactions. For example, *Lactuca serriola* and *Centaurea solstitialis* strongly overlapped in the cool–dry treatment, but in warm–dry conditions, *Lactuca serriola* maintained its flowering phenology while *Centaurea solstitialis* advanced both its onset and end of flowering. These differing responses dramatically reduced the species’ coflowering strength in the warm–dry environment relative to the cool–wet environment (see Supplemental Information). Such examples illustrate how even within a functional group, species-specific sensitivities to warming and drying can erode cohesion within a coflowering module and contribute to broader community-level declines in modularity and coflowering.

### Conclusions and future directions

Network analysis proved a powerful tool for quantifying climate-driven restructuring of community flowering phenology. Although warming and drying produced distinct responses across functional groups, heterogeneity among species within those groups ultimately reduced community-wide coflowering through reduced overlap among summer-flowering species. These results demonstrate that climate-driven shifts in flowering phenology do not necessarily translate into changes in coflowering within functional groups or entire communities. Instead, changes in coflowering likely depend on how climate change differentially affects groups of species flowering at different times of year and the degree of synchrony in phenological responses within these functional groups. Understanding when and how climate change will reorganize coflowering networks therefore requires greater insight into the ecological and evolutionary drivers of species- and functional group-specific phenological responses. Future work should examine the environmental contexts that favor coordinated versus idiosyncratic phenological responses and integrate long-term phenological records, herbarium collections, and community science datasets to assess how coflowering has changed across diverse plant communities over broad spatial and temporal scales.

In addition, we showed that climate-driven shifts in flowering phenology affected seed production, but our experiment could not isolate the fitness consequences of altered coflowering because pollinator communities were shared across treatments and unaffected by our climate manipulations (Greenleaf et al. 2007; Lysenkov et al. 2009). Reduced late-season coflowering may decrease competition for pollinators, heterospecific pollen transfer, and resource competition, potentially increasing plant fitness (Gulmon et al. 1983; Nord and Lynch 2009; Mitchell et al. 2009; Mesgaran et al. 2017; Tiusanen et al. 2020; Cleland and Wolkovich 2024). Conversely, lower floral neighborhood diversity may reduce pollinator attraction (Ghazoul et al. 2006; Liao et al. 2011), while weakened and less diverse coflowering among late-season annuals could diminish nectar resources for pollinators dependent on summertime flowers (Memmott et al. 2007; Rafferty et al. 2014; Ogilvie and Forrest 2017). Determining the net consequences of altered coflowering therefore requires studies that jointly examine plant phenology, pollinator dynamics, and fitness outcomes. Long-term observational studies linking climate-driven changes in coflowering to seed set and pollinator communities, together with experiments manipulating both abiotic conditions and biotic interactions, will be essential for predicting how plant–pollinator communities reorganize under ongoing climate change.

## Acknowledgments

We thank the many people and organizations who made this work possible. We are grateful to the technicians, interns, and volunteers whose efforts supported this project, including Emma Cieslik, Sabrina Deriche, Morgan Harper, Jonathon Howell, Stefany Maldonado, Julie Marco, Perry McCarty, Meera Putz, and Anthony Sifuentes. We also thank Jasper Ridge Biological Preserve (’Ootchamin ’Ooyakma) for providing seeds and Waste Management, Inc. for supplying soils to our experiments. Funding support was provided by Carnegie Science, the California Native Grasslands Association (CNGA) Grassland Research Awards for Student Scholarship (GRASS), and the Irene Brown Fund for Nature-Related Research and Environmental Justice.

