## Supplemental Information for "Warming and precipitation change alter flowering phenology and coflowering networks in California serpentine grasslands"

APPENDIX S1: SUPPLEMENTARY INFORMATION

### Warming and precipitation change alter flowering phenology and coflowering networks in California serpentine grasslands

Andrea N. Nebhut[1] and Jeffrey S. Dukes

SUPPLEMENTARY METHODS

#### Experimental set-up

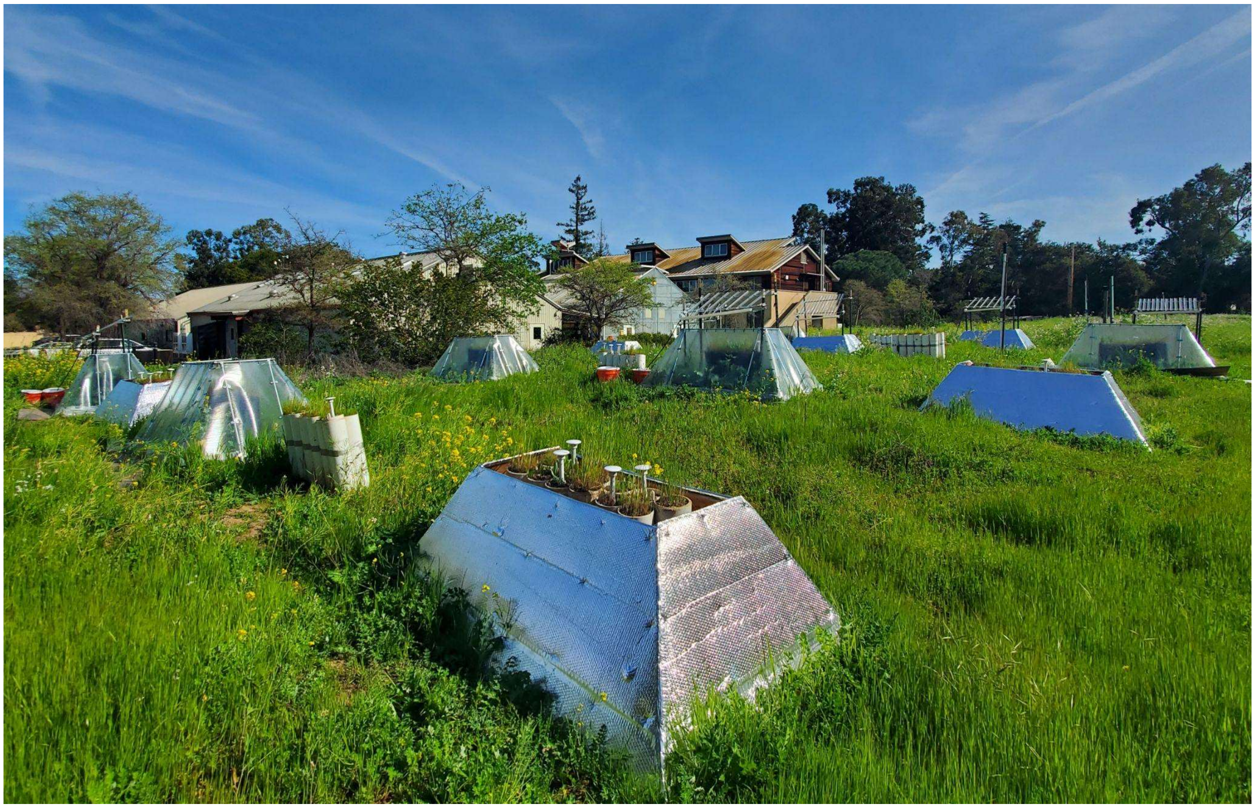

*Figure S1*. Photograph of the experiment.

#### Springtime temperature and soil moisture

We quantified soil surface temperature (°C) and soil moisture (volumetric water content, %) from 19 March to 20 June 2024 in all no-plant control mesocosms and all single functional group mesocosms (i.e., mesocosms containing only early-season natives, early-season non-natives, late-season natives, or late-season non-natives). Measurements were collected using TMS-4 dataloggers (TOMST, Prague, Czech Republic) deployed in the 40%, 100%, and 190% precipitation treatments, for a total of 54 monitored mesocosms (*Fig. S2–S3*).

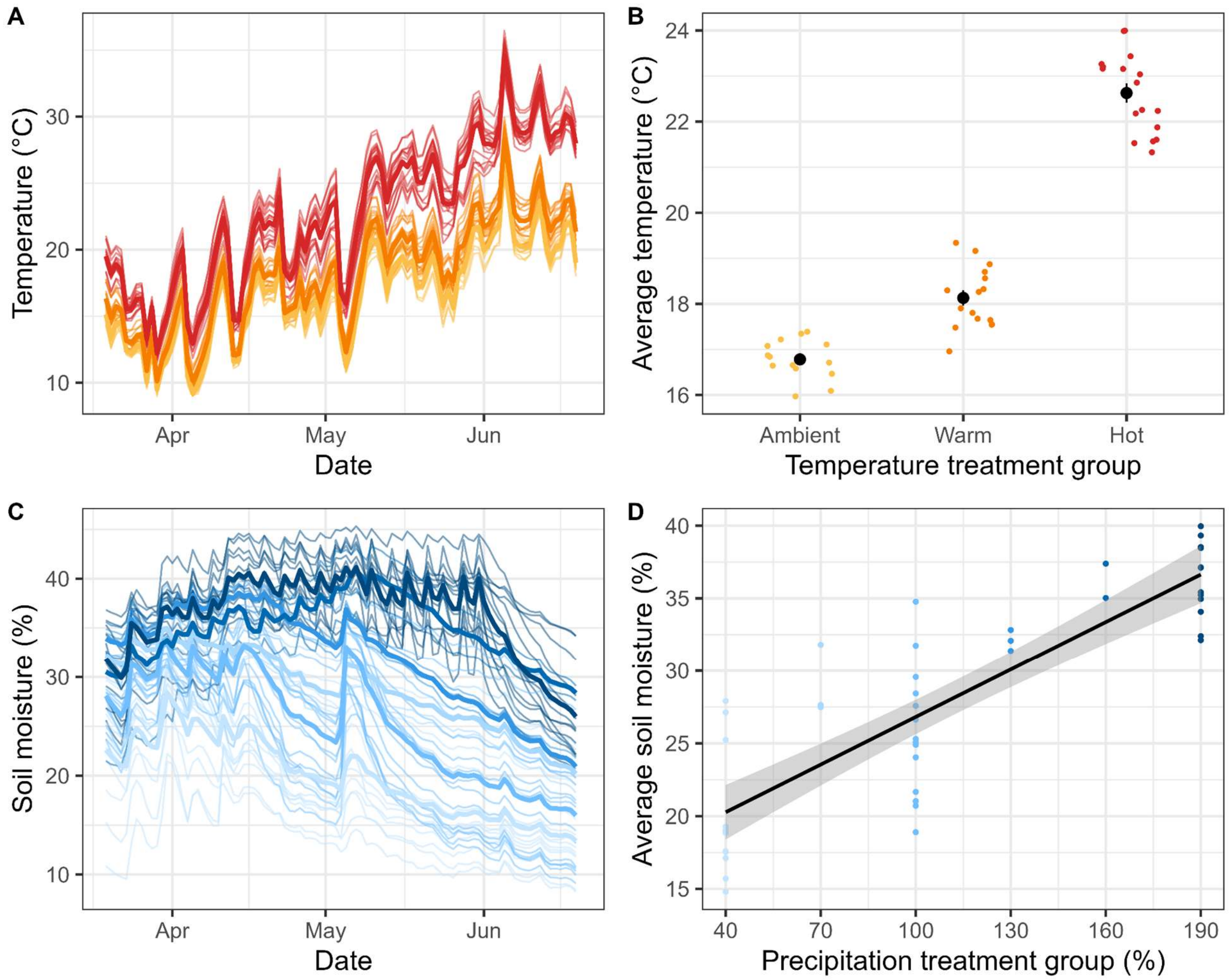


*Figure S2.* Daily and seasonal climate patterns. A) Daily soil surface temperature (°C) and C) soil moisture (%) in each mesocosm over the spring (March 20–June 21, 2024). Thin lines show individual mesocosms; thick lines show treatment group averages. B) Seasonal average soil surface temperature (°C) by climate treatment (ambient, warm, hot). Points represent individual mesocosms; lines show mean ± SE. D) Seasonal average soil moisture (%) by precipitation treatment (% of ambient). Points show individual mesocosms; lines show best fit ± 90% CI.

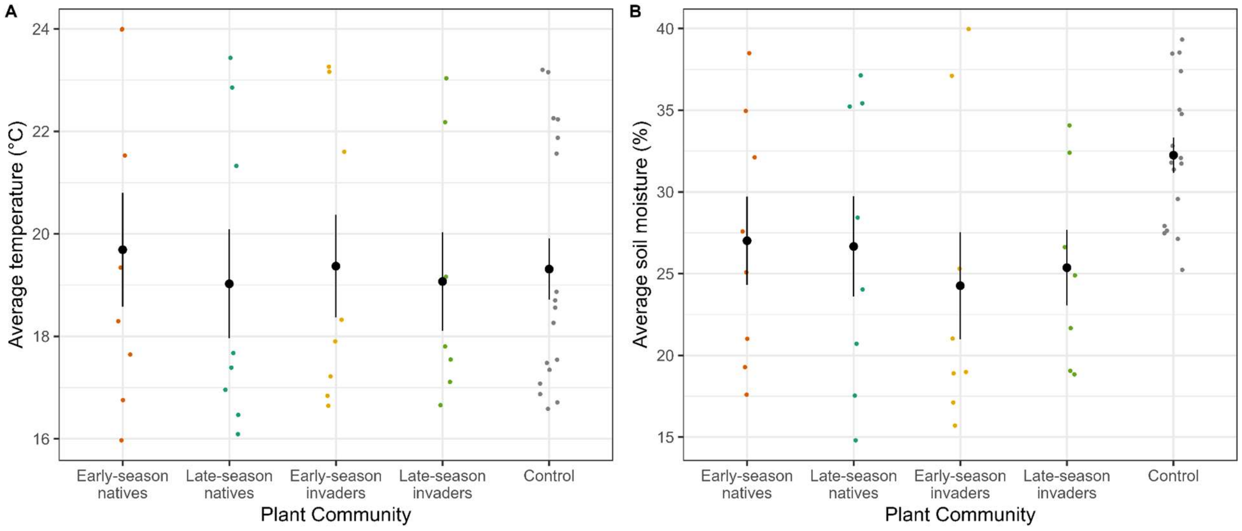


*Figure S3.* A) Seasonal average soil surface temperature (°C) and B) seasonal average soil moisture (%) across community functional compositions. Points represent individual mesocosms; lines show means ± SE.

To characterize springtime climate conditions in the remaining 162 mesocosms, we used a random forest modeling approach implemented in the *randomForest* R package (Liaw and Wiener 2002). The models predicted average soil temperature and moisture over the same measurement period based on the temperature (ambient, warm, and hot) and precipitation (40–190%) treatment groups, as well as the presence of early- or late-season, native or non-native annuals, the block number, and the side (north or south) of the block where the mesocosm was located to account for spatial heterogeneity across our experimental site.

The random forest models adequately predicted the climate conditions experienced by the mesocosms (*Fig. S4*). The temperature model explained 83.2% of the variance (mean squared residual = 1.14). Among predictor variables, temperature treatment group and block identity contributed most strongly to model accuracy, while precipitation, functional group composition, and side had comparatively minor effects. Likewise, the soil moisture model explained 74.1% of the variance (mean squared residual = 13.85). Precipitation was by far the most influential predictor of soil moisture, followed by block identity and, to a lesser extent, the presence of early-season invaders. Ambient temperature and most functional-group indicators contributed relatively little to model performance.

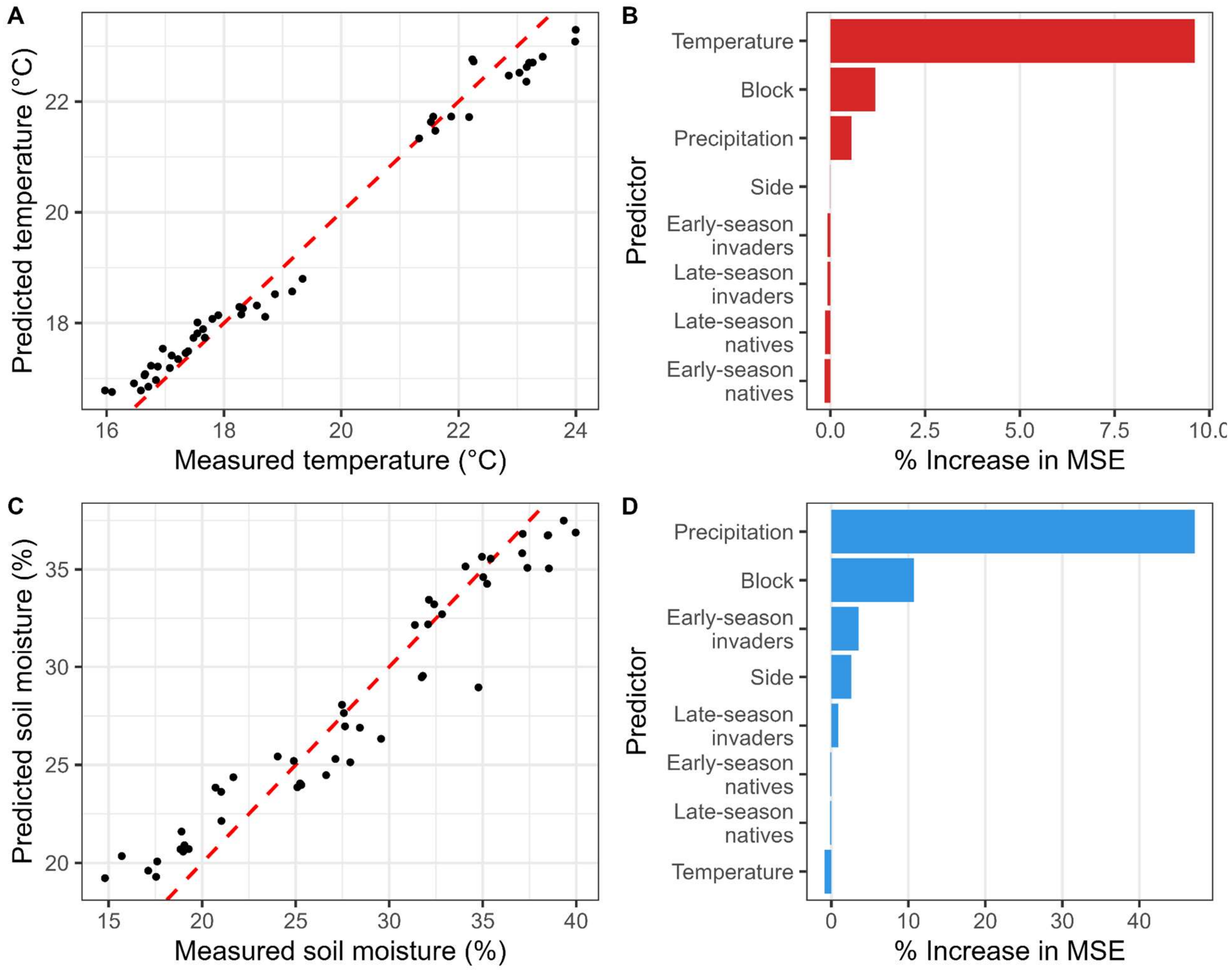


*Figure S4.* Random forest model results. Predicted vs. observed A) soil surface temperature (°C) and C) soil moisture (%) with the 1:1 line, and the variable importance for the B) temperature model and D) soil moisture model, expressed as the percent increase in mean squared error (%IncMSE) when each predictor is permuted, indicating the relative contribution of each predictor to model accuracy.

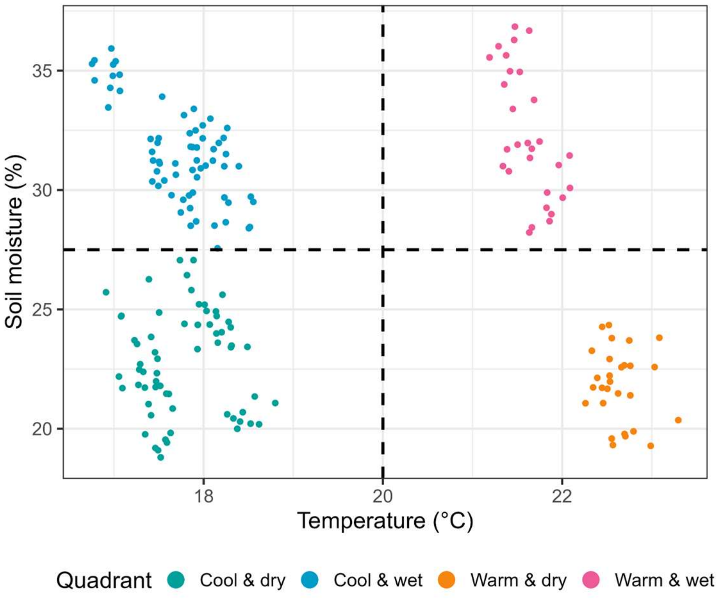


*Figure S5.* Temperature and soil moisture environment of mesocosm communities. Each point represents a single mesocosm community, plotted by its average springtime (March 19–June 20, 2024) soil surface temperature (°C) and soil moisture (%). These mesocosms are grouped into four environmental quadrants based on natural breaks in the data: cool & dry (green), cool & wet (blue), warm & dry (orange), and warm & wet (pink).

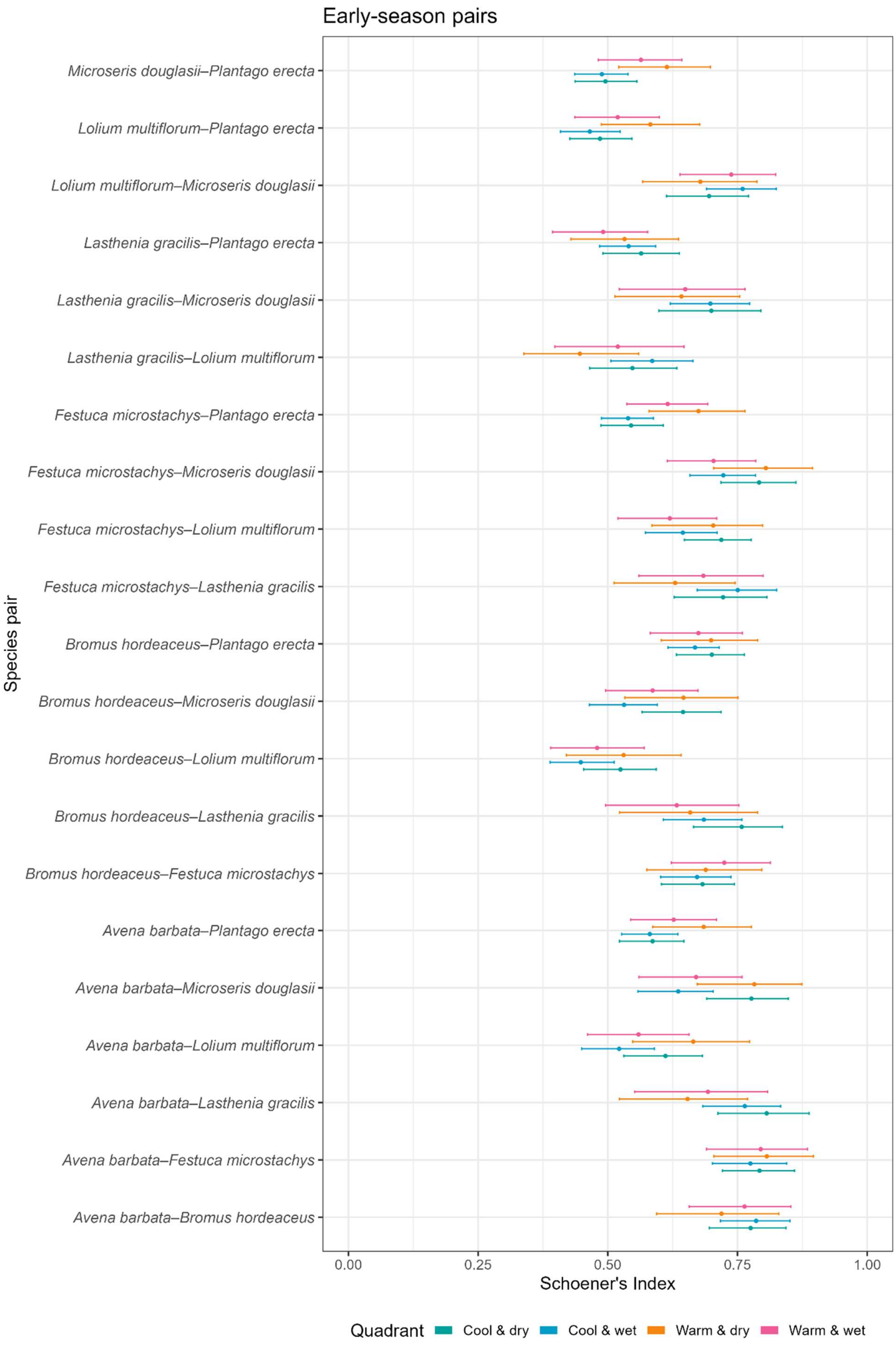
Early-season pairs
Microseris douglasii–Plantago erecta
Lolium multiflorum–Plantago erecta
Lolium multiflorum–Microseris douglasii
Lasthenia gracilis–Plantago erecta
Lasthenia gracilis–Microseris douglasii
Lasthenia gracilis–Lolium multiflorum
Festuca microstachys–Plantago erecta
Festuca microstachys–Microseris douglasii
Festuca microstachys–Lolium multiflorum
Festuca microstachys–Lasthenia gracilis
Bromus hordeaceus–Plantago erecta
Bromus hordeaceus–Microseris douglasii
Bromus hordeaceus–Lolium multiflorum
Bromus hordeaceus–Lasthenia gracilis
Bromus hordeaceus–Festuca microstachys
Avena barbata–Plantago erecta
Avena barbata–Microseris douglasii
Avena barbata–Lolium multiflorum
Avena barbata–Lasthenia gracilis
Avena barbata–Festuca microstachys
Avena barbata–Bromus hordeaceus
Species pair
0.00
0.25
0.50
0.75
1.00
Schoener's Index
Quadrant
Cool & dry
Cool & wet
Warm & dry
Warm & wet

*Figure S6*. Bootstrapped Schoener's index for early-season species pairs. Shown are Schoener's index values (mean ± SE) derived from bootstrap resampling for each species pair. Colors indicate the climate context in which the pair co-occurred: cool–dry (green), cool–wet (blue), warm–dry (orange), and warm–wet (pink).

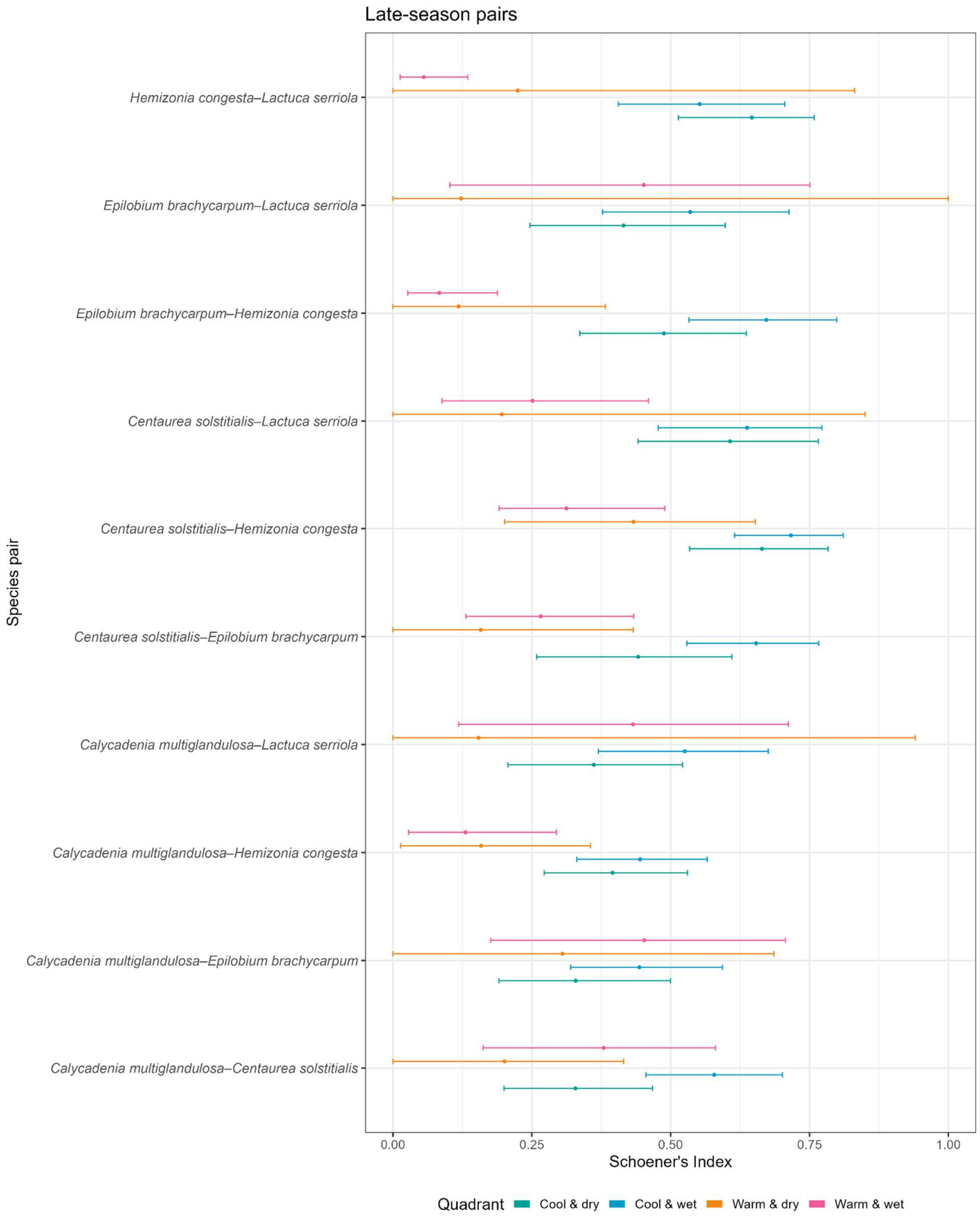


*Figure S7*. Bootstrapped Schoener's index for late-season species pairs. Shown are Schoener's index values (mean ± SE) derived from bootstrap resampling for each species pair. Colors indicate the climate context in which the pair co-occurred: coo–dry (green), cool–wet (blue), warm–dry (orange), and warm–wet (pink).

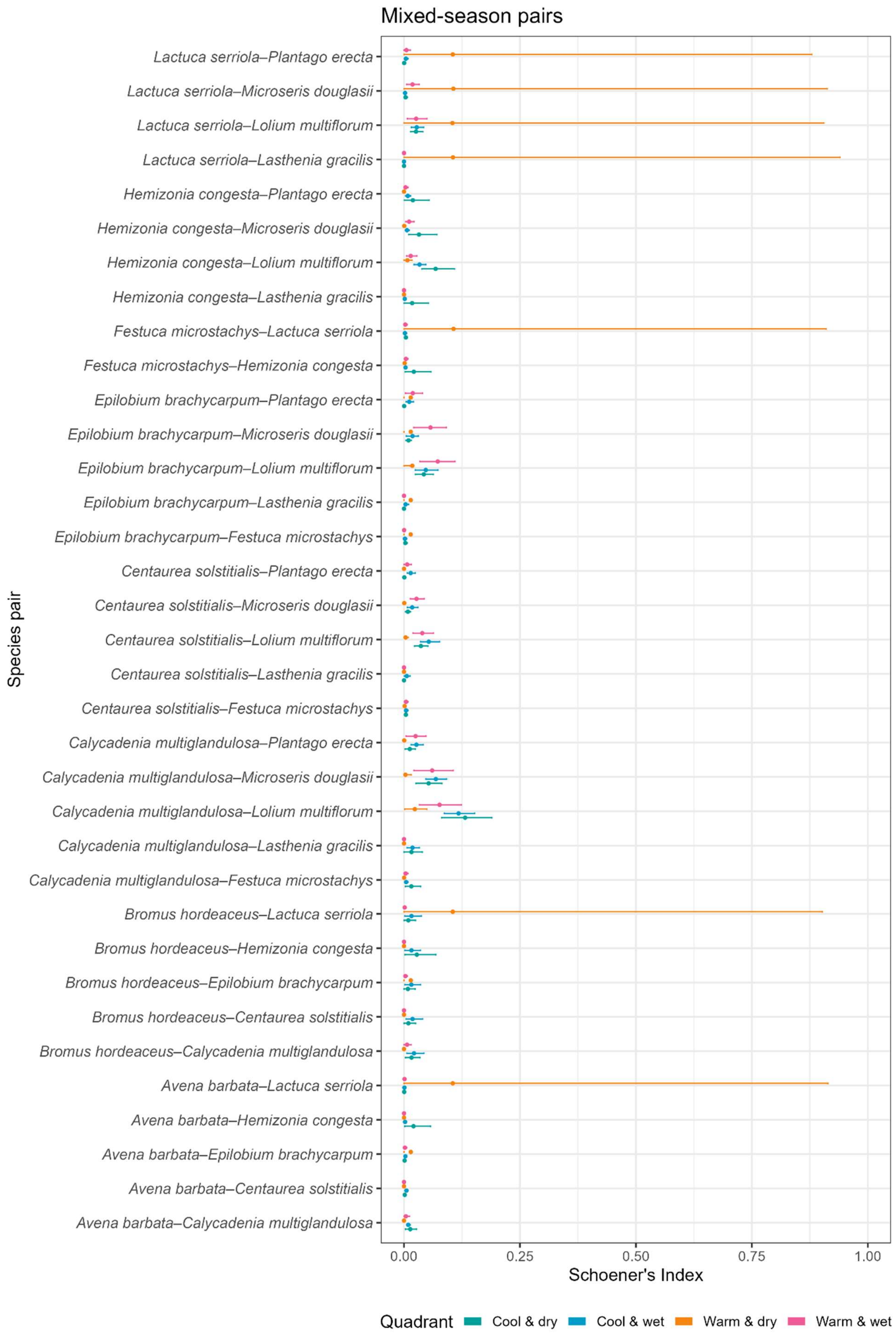
Mixed-season pairs
Lactuca serriola–Plantago erecta
Lactuca serriola–Microseris douglasii
Lactuca serriola–Lolium multiflorum
Lactuca serriola–Lasthenia gracilis
Hemizonia congesta–Plantago erecta
Hemizonia congesta–Microseris douglasii
Hemizonia congesta–Lolium multiflorum
Hemizonia congesta–Lasthenia gracilis
Festuca microstachys–Lactuca serriola
Festuca microstachys–Hemizonia congesta
Epilobium brachycarpum–Plantago erecta
Epilobium brachycarpum–Microseris douglasii
Epilobium brachycarpum–Lolium multiflorum
Epilobium brachycarpum–Lasthenia gracilis
Epilobium brachycarpum–Festuca microstachys
Centaurea solstitialis–Plantago erecta
Centaurea solstitialis–Microseris douglasii
Centaurea solstitialis–Lolium multiflorum
Centaurea solstitialis–Lasthenia gracilis
Centaurea solstitialis–Festuca microstachys
Calycadenia multiglandulosa–Plantago erecta
Calycadenia multiglandulosa–Microseris douglasii
Calycadenia multiglandulosa–Lolium multiflorum
Calycadenia multiglandulosa–Lasthenia gracilis
Calycadenia multiglandulosa–Festuca microstachys
Bromus hordeaceus–Lactuca serriola
Bromus hordeaceus–Hemizonia congesta
Bromus hordeaceus–Epilobium brachycarpum
Bromus hordeaceus–Centaurea solstitialis
Bromus hordeaceus–Calycadenia multiglandulosa
Avena barbata–Lactuca serriola
Avena barbata–Hemizonia congesta
Avena barbata–Epilobium brachycarpum
Avena barbata–Centaurea solstitialis
Avena barbata–Calycadenia multiglandulosa
Species pair
0.00
0.25
0.50
0.75
1.00
Schoener's Index
Quadrant
Cool & dry
Cool & wet
Warm & dry
Warm & wet

*Figure S8*. Bootstrapped Schoener's index for early- and late-season species pairs. Shown are Schoener's index values (mean ± SE) derived from bootstrap resampling for each species pair. Colors indicate the climate context in which the pair co-occurred: coo–dry (green), cool–wet (blue), warm–dry (orange), and warm–wet (pink).

SUPPLEMENTARY RESULTS

*Table S1.* Standardized path coefficients and p-values for all relationships in the early- and late-season annual structural equation models. Both models showed adequate fit to the data, and no modifications to the initial model structure were required.

| Response | Predictor | Early-season annuals | | | Late-season annuals | | |
|---|---|---|---|---|---|---|---|
| | | df | Std. estimate | p | df | Std. estimate | p |
| Biomass | Temperature | 568.01 | -0.043 | 0.67 | 154.34 | 0.265 | 0.45 |
| Biomass | Soil moisture | 568.02 | -0.114 | 0.59 | 154.26 | 0.556 | 0.45 |
| Biomass | Temperature * soil moisture | 568.02 | 0.130 | 0.56 | 154.31 | -0.368 | 0.64 |
| # flowers | Biomass | 550.50 | 0.719 | 0.00 | 155.60 | 0.706 | 0.00 |
| # flowers | Temperature | 567.06 | -0.036 | 0.66 | 153.22 | -0.464 | 0.04 |
| # flowers | Soil moisture | 567.07 | 0.103 | 0.55 | 153.19 | -0.667 | 0.16 |
| # flowers | Temperature * soil moisture | 567.07 | -0.077 | 0.67 | 153.19 | 0.717 | 0.17 |
| Onset of flowering | # flowers | 570.37 | -0.411 | 0.00 | 150.99 | -0.053 | 0.66 |
| Onset of flowering | Biomass | 571.71 | -0.232 | 0.00 | 152.84 | -0.228 | 0.05 |
| Onset of flowering | Temperature | 566.00 | -0.037 | 0.73 | 153.09 | 0.657 | 0.06 |
| Onset of flowering | Soil moisture | 566.02 | 0.293 | 0.20 | 152.70 | 2.157 | 0.00 |
| Onset of flowering | Temperature * soil moisture | 566.01 | -0.302 | 0.21 | 152.68 | -2.205 | 0.01 |

| | | | | | | | |
|---|---|---|---|---|---|---|---|
| Peak of flowering | Onset of flowering | 483.66 | 0.521 | 0.00 | 154.69 | 0.774 | 0.00 |
| Peak of flowering | # flowers | 548.76 | 0.147 | 0.02 | 144.45 | 0.275 | 0.00 |
| Peak of flowering | Biomass | 496.93 | 0.102 | 0.12 | 143.95 | 0.057 | 0.45 |
| Peak of flowering | Temperature | 564.95 | -0.503 | 0.00 | 153.27 | -0.005 | 0.98 |
| Peak of flowering | Soil moisture | 565.16 | -0.597 | 0.02 | 153.29 | 0.142 | 0.77 |
| Peak of flowering | Temperature * soil moisture | 565.11 | 0.769 | 0.00 | 153.17 | -0.185 | 0.73 |
| End of flowering | Onset of flowering | 531.47 | -0.012 | 0.80 | 150.14 | -0.154 | 0.03 |
| End of flowering | Peak of flowering | 570.00 | 0.245 | 0.00 | 153.73 | 0.849 | 0.00 |
| End of flowering | # flowers | 552.82 | 0.152 | 0.01 | 128.35 | 0.194 | 0.01 |
| End of flowering | Biomass | 497.27 | 0.157 | 0.01 | 134.42 | 0.042 | 0.51 |
| End of flowering | Temperature | 564.75 | -0.179 | 0.12 | 152.58 | -0.170 | 0.39 |
| End of flowering | Soil moisture | 564.44 | 0.532 | 0.03 | 152.56 | -0.120 | 0.77 |
| End of flowering | Temperature * soil moisture | 564.43 | -0.250 | 0.32 | 152.42 | 0.103 | 0.82 |
| Duration of flowering | Onset of flowering | 59.53 | -0.525 | 0.00 | 54.23 | -0.731 | 0.00 |

| | | | | | | | |
|---|---|---|---|---|---|---|---|
| Duration of flowering | End of flowering | 294.45 | 0.917 | 0.00 | 63.51 | 1.121 | 0.00 |
| Duration of flowering | # flowers | 135.18 | -0.001 | 0.01 | 15.33 | -0.002 | 0.13 |
| Duration of flowering | Biomass | 77.87 | 0.001 | 0.03 | 38.21 | -0.002 | 0.16 |
| Duration of flowering | Temperature | 564.74 | 0.000 | 0.58 | 153.49 | 0.000 | 0.93 |
| Duration of flowering | Soil moisture | 564.51 | -0.001 | 0.61 | 153.60 | -0.002 | 0.79 |
| Duration of flowering | Temperature * soil moisture | 563.68 | 0.001 | 0.63 | 153.71 | 0.002 | 0.83 |
| # seeds | Onset of flowering | 562.23 | -13.496 | 0.00 | 148.93 | -9.336 | 0.01 |
| # seeds | Peak of flowering | 562.71 | 0.063 | 0.02 | 150.04 | -0.093 | 0.44 |
| # seeds | End of flowering | 562.25 | 23.411 | 0.00 | 148.87 | 14.484 | 0.01 |
| # seeds | Duration of flowering | 562.25 | -25.373 | 0.00 | 148.83 | -12.779 | 0.01 |
| # seeds | # flowers | 564.64 | 0.549 | 0.00 | 121.29 | 0.586 | 0.00 |
| # seeds | Biomass | 566.35 | 0.587 | 0.00 | 136.59 | 0.228 | 0.01 |
| # seeds | Temperature | 562.03 | -0.131 | 0.09 | 150.25 | -0.250 | 0.31 |
| # seeds | Soil moisture | 561.97 | -0.380 | 0.02 | 150.31 | -0.559 | 0.28 |

| | | | | | | | |
|---|---|---|---|---|---|---|---|
| # seeds | Temperature * soil moisture | 561.97 | 0.403 | 0.02 | 150.18 | 0.626 | 0.27 |

*Table S2*. Statistical results of the LMMs investigating the relationship between the species, climate quadrant, and their interaction on the onset, peak, end, and duration of flowering, the maximum number of flowers observed, and the number of seeds produced.

| | | Onset | Peak | End | Duration | Maximum # flowers | # seeds |
|---|---|---|---|---|---|---|---|
| Species | $df_{num}$ | 11 | 11 | 11 | 11 | 11 | 11 |
| | $df_{denom}$ | 684.996 | 688.750 | 681.709 | 684.188 | 685.220 | 655.110 |
| | F | 449.148 | 396.283 | 160.141 | 40.144 | 122.430 | 46.701 |
| | p | < 0.001 | < 0.001 | < 0.001 | < 0.001 | < 0.001 | < 0.001 |
| Climate treatment | $df_{num}$ | 3 | 3 | 3 | 3 | 3 | 3 |
| | $df_{denom}$ | 29.502 | 30.550 | 25.032 | 30.406 | 282.330 | 295.570 |
| | F | 22.886 | 32.605 | 38.246 | 17.117 | 3.493 | 2.896 |
| | p | < 0.001 | < 0.001 | < 0.001 | < 0.001 | 0.02 | 0.04 |
| Species * Climate treatment | $df_{num}$ | 33 | 33 | 33 | 33 | 33 | 33 |
| | $df_{denom}$ | 684.766 | 689.860 | 680.938 | 683.446 | 683.710 | 654.770 |
| | F | 3.371 | 2.911 | 2.922 | 2.302 | 2.153 | 1.951 |
| | p | < 0.001 | < 0.001 | < 0.001 | < 0.001 | < 0.001 | 0.001 |

*Table S3*. Estimated marginal means and standard errors of the onset, peak, end, and duration of flowering, maximum number of flowers observed, and number of seeds produced for each species in each of the four climate quadrats. Significance determinations derived from post-hoc Tukey's tests; species with the same uppercase letters are not significantly different from each other, and climate treatments within a species with the same lowercase letters are not significantly different from each other.

| Species | Climate treatment | Onset (DOY) | | | Peak (DOY) | | | End (DOY) | | | Duration (# days) | | | Max. # flowers | | | # seeds | | |
|---|---|---|---|---|---|---|---|---|---|---|---|---|---|---|---|---|---|---|---|
| | | Mean | SE | Group | Mean | SE | Group | Mean | SE | Group | Mean | SE | Group | Mean | SE | Group | Mean | SE | Group |
| *Plantago erecta* | All | 64.8 | 1.3 | A | 96.1 | 1.3 | AB | 130.7 | 2.0 | B | 65.9 | 2.0 | E | 10.8 | 1.1 | C | 86.6 | 1.1 | DE |
| | Cool & dry | 64.1 | 2.3 | ab | 95.5 | 2.3 | a | 123.9 | 3.4 | a | 59.6 | 3.5 | ab | 11.0 | 1.1 | ab | 106.5 | 1.2 | b |
| | Cool & wet | 59.1 | 2.1 | a | 95.9 | 2.1 | a | 148.3 | 3.1 | b | 89.5 | 3.2 | c | 16.9 | 1.1 | b | 176.5 | 1.2 | b |
| | Warm & dry | 72.2 | 3.1 | b | 95.4 | 3.1 | a | 119.0 | 4.6 | a | 46.8 | 4.7 | a | 7.7 | 1.2 | a | 37.7 | 1.3 | a |
| | Warm & wet | 63.9 | 3.1 | ab | 97.5 | 3.1 | a | 131.7 | 4.6 | a | 67.8 | 4.7 | b | 9.5 | 1.2 | a | 79.4 | 1.3 | ab |
| *Bromus hordeaceus* | All | 74.7 | 1.6 | B | 98.8 | 1.6 | AB | 129.5 | 2.4 | AB | 54.8 | 2.5 | D | 26.6 | 1.1 | E | 154.2 | 1.1 | EF |
| | Cool & dry | 75.1 | 2.6 | a | 104.3 | 2.7 | a | 130.9 | 3.9 | ab | 55.6 | 4.1 | ab | 26.4 | 1.2 | a | 157.1 | 1.3 | a |
| | Cool & wet | 75.8 | 2.6 | a | 101.1 | 2.7 | a | 140.4 | 3.9 | b | 64.9 | 4.1 | b | 26.6 | 1.2 | a | 160.5 | 1.3 | a |
| | Warm & dry | 75.2 | 3.7 | a | 92.7 | 3.8 | a | 116.1 | 5.5 | a | 40.9 | 5.7 | a | 22.4 | 1.2 | a | 92.9 | 1.4 | a |
| | Warm & wet | 72.7 | 3.7 | a | 97.4 | 3.8 | a | 130.4 | 5.5 | ab | 57.7 | 5.7 | ab | 31.9 | 1.2 | a | 241.5 | 1.4 | a |
| *Avena barbata* | All | 79.4 | 1.6 | BC | 104.1 | 1.6 | BC | 135.3 | 2.4 | B | 55.9 | 2.5 | DE | 31.4 | 1.1 | EF | 45.9 | 1.1 | C |
| | Cool & dry | 82.6 | 2.6 | a | 107.6 | 2.7 | a | 137.3 | 3.9 | ab | 54.5 | 4.1 | a | 31.9 | 1.2 | a | 45.5 | 1.3 | a |
| | Cool & wet | 78.7 | 2.6 | a | 106.4 | 2.7 | a | 151.2 | 3.9 | b | 72.8 | 4.1 | b | 38.4 | 1.2 | a | 61.9 | 1.3 | a |
| | Warm & dry | 79.3 | 3.7 | a | 100.7 | 3.8 | a | 121.5 | 5.5 | a | 42.3 | 5.7 | a | 24.2 | 1.2 | a | 32.9 | 1.4 | a |
| | Warm & wet | 76.9 | 3.7 | a | 101.7 | 3.8 | a | 131.0 | 5.5 | a | 54.1 | 5.7 | a | 32.8 | 1.2 | a | 48.0 | 1.4 | a |
| *Loli* | All | 80.5 | 1.6 | BC | 120.9 | 1.6 | E | 164.7 | 2.4 | D | 84.2 | 2.5 | F | 103.9 | 1.1 | G | 461.3 | 1.1 | G |

| | | | | | | | | | | | | | | | | | | | |
|---|---|---|---|---|---|---|---|---|---|---|---|---|---|---|---|---|---|---|---|
| | Cool & dry | 81.2 | 2.6 | a | 121.6 | 2.7 | ab | 173.5 | 3.9 | b | 92.1 | 4.1 | b | 86.6 | 1.2 | a | 337.5 | 1.3 | a |
| | Cool & wet | 84.3 | 2.6 | a | 127.6 | 2.7 | b | 181.2 | 3.9 | b | 97.2 | 4.1 | b | 84.1 | 1.2 | a | 350.5 | 1.3 | a |
| | Warm & dry | 78.3 | 3.7 | a | 112.5 | 3.8 | a | 136.1 | 5.5 | a | 57.8 | 5.7 | a | 113.7 | 1.2 | a | 475.2 | 1.4 | a |
| | Warm & wet | 78.0 | 3.7 | a | 121.8 | 3.8 | ab | 167.8 | 5.5 | b | 89.7 | 5.7 | b | 140.6 | 1.2 | a | 805.6 | 1.4 | a |
| *Festuca microstachys* | All | 81.8 | 1.3 | C | 109.2 | 1.3 | CD | 132.1 | 2.0 | B | 50.3 | 2.0 | CD | 44.8 | 1.1 | F | 266.9 | 1.1 | FG |
| | Cool & dry | 86.1 | 2.2 | ab | 113.5 | 2.3 | b | 135.6 | 3.3 | b | 49.3 | 3.4 | a | 41.6 | 1.1 | a | 224.2 | 1.2 | a |
| | Cool & wet | 86.9 | 2.1 | b | 114.3 | 2.1 | b | 138.8 | 3.1 | b | 52.2 | 3.2 | a | 40.3 | 1.1 | a | 230.6 | 1.2 | a |
| | Warm & dry | 77.0 | 3.1 | a | 101.5 | 3.1 | a | 120.5 | 4.6 | a | 43.5 | 4.7 | a | 50.0 | 1.2 | a | 320.5 | 1.3 | a |
| | Warm & wet | 77.4 | 3.1 | ab | 107.5 | 3.1 | ab | 133.6 | 4.6 | ab | 56.2 | 4.7 | a | 48.0 | 1.2 | a | 306.3 | 1.3 | a |
| *Microseris douglasii* | All | 88.9 | 1.4 | D | 111.2 | 1.4 | D | 146.7 | 2.0 | C | 57.8 | 2.1 | DE | 15.6 | 1.1 | BC | 224.9 | 1.1 | F |
| | Cool & dry | 91.9 | 2.3 | ab | 113.5 | 2.4 | b | 149.0 | 3.5 | b | 56.8 | 3.6 | b | 18.7 | 1.1 | a | 174.9 | 1.2 | a |
| | Cool & wet | 95.4 | 2.2 | b | 119.3 | 2.3 | b | 170.1 | 3.3 | c | 75.0 | 3.4 | c | 16.1 | 1.1 | a | 178.8 | 1.2 | a |
| | Warm & dry | 83.1 | 3.1 | a | 100.4 | 3.1 | a | 118.6 | 4.6 | a | 35.4 | 4.7 | a | 14.0 | 1.2 | a | 258.9 | 1.3 | a |
| | Warm & wet | 85.2 | 3.1 | a | 111.6 | 3.1 | ab | 149.1 | 4.6 | b | 63.9 | 4.7 | bc | 14.0 | 1.2 | a | 315.8 | 1.3 | a |
| *Lasthenia gracilis* | All | 93.4 | 1.7 | D | 104.3 | 1.7 | BCD | 119.5 | 2.5 | A | 26.2 | 2.6 | AB | 2.9 | 1.1 | A | 30.7 | 1.2 | BC |
| | Cool & dry | 92.7 | 2.7 | ab | 104.5 | 2.7 | ab | 120.3 | 4.0 | b | 27.4 | 4.1 | ab | 3.5 | 1.2 | a | 45.7 | 1.3 | a |
| | Cool & wet | 99.6 | 2.5 | b | 112.6 | 2.5 | b | 140.1 | 3.6 | c | 41.1 | 3.8 | b | 3.6 | 1.1 | a | 39.0 | 1.3 | a |
| | Warm & dry | 85.0 | 3.4 | a | 95.1 | 3.4 | a | 102.9 | 5.0 | a | 18.0 | 5.1 | a | 2.7 | 1.2 | a | 29.4 | 1.4 | a |
| | Warm & wet | 96.3 | 4.6 | ab | 104.7 | 4.6 | ab | 114.5 | 6.6 | ab | 18.3 | 7.0 | a | 2.2 | 1.3 | a | 17.0 | 1.7 | a |
| *Calycadenia multiglandulosa* | All | 164.7 | 3.3 | E | 184.1 | 3.3 | F | 196.1 | 4.7 | E | 31.7 | 5.1 | AB | 5.4 | 1.2 | AB | 8.5 | 1.4 | A |
| | Cool & dry | 182.1 | 4.9 | b | 196.2 | 4.9 | b | 208.7 | 6.9 | b | 26.4 | 7.5 | a | 3.4 | 1.3 | a | 8.0 | 1.7 | a |
| | Cool & wet | 176.5 | 3.7 | b | 200.8 | 3.7 | b | 218.6 | 5.3 | b | 42.8 | 5.7 | a | 8.9 | 1.2 | b | 10.3 | 1.5 | a |
| | Warm & dry | 149.8 | 7.4 | a | 159.2 | 7.4 | a | 159.7 | 10.5 | a | 10.8 | 11.4 | a | 3.4 | 1.5 | ab | 4.7 | 1.9 | a |
| | Warm & wet | 150.4 | 9.0 | a | 180.3 | 9.1 | ab | 197.5 | 12.7 | ab | 46.8 | 13.9 | a | 8.0 | 1.7 | ab | 13.2 | 2.2 | a |
| *Centau rea solstiti alis* | All | 173.6 | 3.0 | EF | 199.5 | 3.0 | G | 228.2 | 4.2 | G | 54.4 | 4.6 | CDE | 2.5 | 1.2 | A | 18.0 | 1.3 | ABC |
| | Cool & dry | 181.6 | 3.3 | bc | 204.9 | 3.4 | ab | 236.6 | 4.8 | ab | 55.1 | 5.1 | a | 2.8 | 1.2 | a | 19.9 | 1.3 | a |
| | Cool & wet | 191.6 | 3.7 | c | 209.0 | 3.7 | b | 243.1 | 5.3 | b | 51.6 | 5.7 | a | 3.2 | 1.2 | a | 38.1 | 1.4 | a |

| | | | | | | | | | | | | | | | | | | | |
|---|---|---|---|---|---|---|---|---|---|---|---|---|---|---|---|---|---|---|---|
| | Warm & dry | 154.8 | 9.0 | a | 194.1 | 9.1 | ab | 204.3 | 12.8 | a | 49.2 | 14.0 | a | 1.0 | 1.7 | a | 5.7 | 2.2 | a |
| | Warm & wet | 166.3 | 5.7 | ab | 189.9 | 5.8 | a | 228.8 | 8.2 | ab | 61.5 | 8.8 | a | 4.1 | 1.4 | a | 23.9 | 1.6 | a |
| *Epilobium brachycarpum* | All | 175.1 | 3.8 | EF | 182.7 | 3.8 | F | 188.1 | 5.3 | E | 13.5 | 5.8 | A | 3.1 | 1.2 | A | 9.5 | 1.4 | AB |
| | Cool & dry | 189.0 | 4.3 | b | 194.8 | 4.3 | b | 203.7 | 6.1 | bc | 14.4 | 6.6 | a | 1.7 | 1.3 | a | 5.3 | 1.4 | a |
| | Cool & wet | 199.9 | 3.3 | b | 210.6 | 3.3 | c | 217.7 | 4.8 | c | 18.3 | 5.1 | a | 4.4 | 1.2 | b | 15.8 | 1.3 | a |
| | Warm & dry | 145.4 | 12.8 | a | 154.2 | 12.8 | a | 152.2 | 18.0 | a | 8.2 | 19.8 | a | 4.3 | 2.0 | ab | 7.9 | 3.0 | a |
| | Warm & wet | 166.3 | 5.7 | a | 171.2 | 5.8 | a | 178.6 | 8.2 | ab | 13.0 | 8.8 | a | 2.9 | 1.4 | ab | 12.4 | 1.6 | a |
| *Hemizonia congesta* | All | 184.7 | 2.1 | F | 208.0 | 2.1 | G | 222.8 | 3.0 | FG | 38.4 | 3.2 | BC | 7.9 | 1.1 | BC | 44.3 | 1.2 | CD |
| | Cool & dry | 189.1 | 2.6 | ab | 215.9 | 2.7 | b | 239.0 | 3.9 | b | 49.5 | 4.1 | bc | 10.2 | 1.2 | b | 47.7 | 1.3 | ab |
| | Cool & wet | 192.2 | 2.5 | b | 223.3 | 2.5 | b | 245.7 | 3.7 | b | 54.1 | 3.8 | c | 13.4 | 1.1 | b | 95.4 | 1.2 | b |
| | Warm & dry | 182.0 | 5.8 | ab | 196.0 | 5.8 | a | 199.0 | 8.2 | a | 18.1 | 8.9 | a | 3.6 | 1.4 | a | 16.2 | 1.6 | a |
| | Warm & wet | 175.6 | 4.9 | a | 196.9 | 4.9 | a | 207.4 | 7.0 | a | 32.0 | 7.5 | ab | 7.7 | 1.3 | ab | 52.1 | 1.5 | ab |
| *Lactuca serriola* | All | 186.7 | 3.9 | F | 195.2 | 3.9 | FG | 206.0 | 5.6 | EF | 19.1 | 6.1 | AB | 30.6 | 1.2 | DEF | 43.4 | 1.4 | BCD |
| | Cool & dry | 194.9 | 4.5 | b | 207.3 | 4.6 | b | 228.8 | 6.5 | b | 34.5 | 7.0 | a | 25.3 | 1.3 | a | 35.8 | 1.5 | a |
| | Cool & wet | 196.8 | 4.5 | b | 204.1 | 4.6 | b | 215.5 | 6.5 | b | 19.0 | 7.0 | a | 17.2 | 1.3 | a | 14.6 | 1.5 | a |
| | Warm & dry | 182.8 | 12.8 | ab | 188.0 | 12.8 | ab | 194.4 | 18.1 | ab | 10.1 | 19.8 | a | 45.0 | 2.1 | a | 81.8 | 3.0 | a |
| | Warm & wet | 172.4 | 6.4 | a | 181.3 | 6.4 | a | 185.3 | 9.1 | a | 12.9 | 9.9 | a | 44.9 | 1.4 | a | 82.7 | 1.7 | a |
